# Recurring infection with ecologically distinct human papillomavirus (HPV) types explains high prevalence and diversity

**DOI:** 10.1101/179341

**Authors:** Sylvia Ranjeva, Edward B. Baskerville, Vanja Dukic, Luisa Villa, Eduardo Lazcano-Ponce, Anna Giuliano, Greg Dwyer, Sarah Cobey

## Abstract

The high prevalence of human papillomavirus (HPV), the most common sexually transmitted infection, arises from the coexistence of over 200 genetically distinct types. Accurately predicting the impact of vaccines that target multiple types requires understanding the factors that determine HPV diversity. The diversity of many pathogens is driven by type-specific or “homologous” immunity, which promotes the spread of variants to which hosts have little immunity. To test for homologous immunity and to identify mechanisms determining HPV transmission, we fitted nonlinear mechanistic models to longitudinal data on genital infections in unvaccinated men. Our results provide no evidence for homologous immunity, instead showing that infection with one HPV type strongly increases the risk of infection with that type for years afterwards. For HPV16, the type responsible for most HPV-related cancers, an initial infection increases the one-year probability of reinfection by 20-fold, and the probability of reinfection remains 14-fold higher two years later. This increased risk occurs in both sexually active and celibate men, suggesting that it arises from auto-inoculation, episodic reactivation of latent virus, or both. Overall our results show that high HPV prevalence and diversity can be explained by a combination of a lack of homologous immunity, frequent reinfections, weak competition between types, and variation in type fitness between host subpopulations. Due to the high risk of reinfection, vaccinating boys that have not yet been exposed may be crucial to reduce prevalence, but our results suggest that there may also be large benefits from vaccinating previously infected individuals.

## Introduction

Human papillomavirus (HPV), a major cause of genital warts and anogenital and oropharyngeal cancers [46, 32, 118], is the most common sexually transmitted infection [119]. While genital HPV infects approximately 40% of women and 45% of men in the USA, over 200 HPV types have been identified, and the prevalence of individual types never exceeds 10% [44]. The quadrivalent vaccine has been effective against the HPV types that cause the greatest burden of disease [29, 70, 115], and a recent nine-valent vaccine covers additional oncogenic types [52]. Predicting the effects of multivalent vaccines on HPV prevalence, however, is difficult without defining the factors that underlie HPV transmission and diversity, which are poorly understood.

Epidemiological theory has shown that the abundance of many pathogens depends on the dynamics of competition for susceptible hosts [66, 112, 18]. Pathogen strains compete by inducing adaptive immune responses that are specific to shared antigens, limiting the growth rates of antigenically similar competitors [18]. The accumulation of specific immunity in the host population to common strains decreases the transmission of those strains and promotes the spread of rarer antigenic variants, a phenomenon known as frequency-dependent selection. Strains that vary antigenically are often organized into groups that vary in dominance in space and time, yielding complex patterns of coexistence [43, 66]. Such immune-mediated competition can partly explain the antigenic and genetic diversity of influenza, pneumococcus, rotavirus, norovirus, *Neisseria meningitidis*, malaria, hepatitis C, HIV, trypanosomes, and other common pathogens [71, 18, 66, 65, 112].

It is unclear how HPV interacts with the immune system during infection, but in principle, distinctions between HPV types may arise due to acquired immunity from B cells and T cells. HPV types are defined by a 10% threshold of dissimilarity in the L1 gene, which codes for the major capsid protein [19]. The outer capsid modulates viral entry into host cells at the epithelial basement membrane [13], and the humoral response to infection is mainly type-specific anti-L1 antibodies [92, 108]. Studies of HPV in T-cell-deficient people show that cellular immunity is important in the control of infection [103, 92, 6, 93, 31], but it remains unclear how cellular immunity achieves type-specific recognition, if at all. In individuals with cancerous lesions from HPV16, cytotoxic T lymphocytes specific to the E6 and E7 oncoproteins are correlated with reduced disease [82, 95]. The specificity of the T cell repertoire to other genes or to the majority of HPV types, however, is not well established.

Efforts to understand the effects of immunity on HPV dynamics must begin with homologous immunity, or protection against repeat infection with the same HPV type. Homologous immunity would limit the prevalence of any type through negative frequency dependence. The traditional assumption is that most HPV-infected individuals permanently clear infection after 1âĂŞ2 years [107, 100, 78], suggesting protective homologous immunity. In this scenario, the elevated cancer risk associated with particular HPV types results from a small fraction of persistent infections [78]. Longitudinal studies, however, have shown that hosts can be infected repeatedly [104, 28, 29]. Although there is some evidence that type-specific antibodies might provide modest protection against future infection in women [6, 28], serum antibody is not a marker of immune protection in men [90, 67]. Thus, the form and strength of homologous immunity remain unclear.

Evidence for competition between HPV types is also conflicting. Immune-mediated competition has been invoked to explain small increases in the prevalence of non-vaccine HPV types following HPV vaccination [53], and to explain weak cross-protection from the vaccine against related non-vaccine types [121]. Mathematical models show that type competition is consistent with observed patterns of HPV prevalence [25, 81]. Nevertheless, there is little empirical evidence for inter-type competition [64, 98], as shown in part by elevated rates of multiple-type compared to single-type infections and frequent concurrent acquisition of HPV types [15, 114]. Type competition may therefore be absent or weak [99, 16].

Meanwhile, the risk of HPV infection appears to depend on differences in demographic and behavioral risk factors between host subpopulations. For example, a host’s number of sexual partners strongly affect infection risk [122, 33, 36, 79, 78], and some evidence suggests that oncogenic and non-oncogenic types are affected differently by variation in numbers of partners among hosts [60, 111]. However, detailed comparisons of risk factors for infection with each HPV type have not been performed. Different risk factors would suggest differences in transmission rates between types among host subpopulations, defining ecological distinctions that could potentially explain type prevalence.

To investigate the factors that determine HPV prevalence and diversity, we fitted mechanistic models of HPV dynamics to an extensive longitudinal dataset. Mechanistic models have long been used in infectious disease ecology to quantify the biological processes that underly pathogen dynamics, but HPV has received comparatively little attention [100]. Mechanistic models of HPV have generally focused on qualitative dynamics [76, 80, 12, 96, 89, 25, 57, 8, 4, 14, 117, 50, 101] and on predictions for health and economic policy [8, 105, 56, 24, 11, 116, 12], relying on informal methods to estimate parameters. Detailed longitudinal studies, however, present an opportunity to use mechanistic models to rigorously test hypotheses using robust statistical methods. Here we use such methods to show that the prevalence and diversity of HPV types are best explained by the combined effects of a lack of competition, whether within or between types, high rates of reinfection or persistence within individuals, and modest differences in high-risk subpopulations between types.

## Results

### Low-prevalence HPV types coexist over time

We fitted models to data from the HPV in Men (HIM) study, which tracks genital HPV infection and demographic and behavioral traits in unvaccinated men sampled at six month intervals over five years [34, 35, 33]. These data show that the mean prevalence of all HPV is 65% but that no type has a mean prevalence greater than 10% (Figure 1A), and that the prevalence of individual types is roughly constant over time (Figure 1B), demonstrating coexistence. The rank prevalence of HPV types is also constant among geographical locations, although absolute prevalence is higher in Brazil than in the USA or Mexico (Figure S3). We analyzed the five HPV types with the highest mean prevalences: HPV62, HPV84, HPV89, HPV16, and HPV51. We also analyzed HPV6, the ninth most prevalent type, because of its clinical significance. HPV6 is included in the quadrivalent vaccine and is one of two HPV types that are together associated with 90% of genital warts [45]. In our models, we accounted for risk factors that have been previously identified as being associated with the incidence and duration of HPV infection in the HIM dataset [34, 33, 86, 84, 94]. The risk factors for any HPV type include markers of increased exposure to infected sexual partners, which is not surprising for a sexually transmitted infection. For example, higher numbers of recent and lifetime sexual partners, either male or female, increase risk, whereas consistent condom use reduces risk among nonmonogamous men. In addition, risk factors include non-sexual behaviors, such as tobacco use, and demographic traits. For example, incidence is lower among ethnic Asians than other ethnic groups, and among individuals having completed education beyond secondary school. Several risk factors, notably circumcision and sexual orientation, differ in their effects among HPV types [1, 85]. Previous work thus highlighted the breadth of risk factors that affect HPV transmission, and type-specific differences in some of these risk factors would reflect meaningful ecological distinctions between HPV types. We therefore included the effects of a diverse set of risk factors in our models.

**Figure 1:**
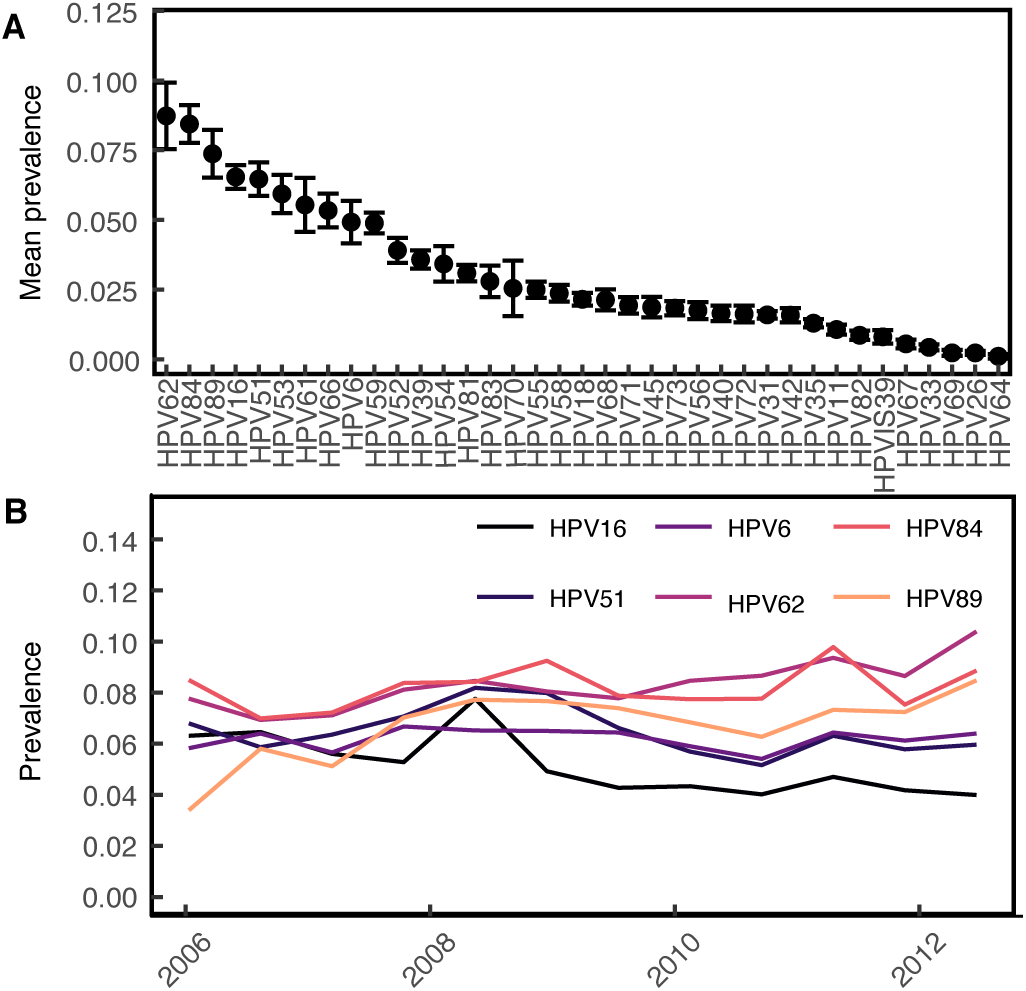
Prevalence of HPV types in the study population. **A** Average prevalence of each HPV type, in order of decreasing prevalence, with standard errors calculated across visits. **B** Prevalence over time of the six HPV types included in our analyses.

### Past infection with HPV confers minimal protection against infection with the same type

Our models test three hypotheses about the dynamics of HPV infection. Under the simplest, or “memoryless” model, the risk of infection depends only on the effects of host- and HPV-type-specific risk factors (Equation 1), with no consideration of immunity. Two more complex models account for the effects of previous infection with the same type. In all models, we assumed that the parameters describing the distribution of the duration of carriage are fixed and type-specific. Our three models then differ only in their assumptions about the instantaneous per-capita infection risk, or “force of infection” [54]. The force of infection for host *i* with HPV type *j* at time *t* is given by *λ*_*ijt*_:

(i) In the memoryless model, risk of infection with type *j* depends only on host specific risk factors and the baseline force of infection for that type, 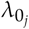:

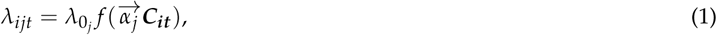 The vector 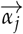 scales the effect of each of the *M* covariates on the force of infection, where ***C*_*it*_** is the covariate matrix. The effects of the covariates on the force of infection are represented by 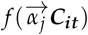, according to:

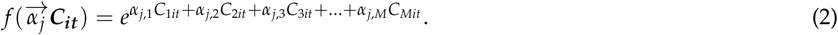
(ii) In the homologous immunity model, protective immunity reduces the probability of reinfection (*p*_subsequent_) relative to the probability of an initial infection (*p*_first_) with the same type:

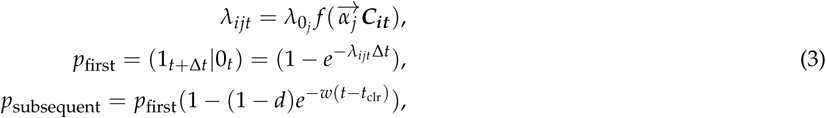 The strength of homologous immunity *d* is constrained to fall between 0 and 1, so that *d* reduces the probability of subsequent infection over time interval Δt. After the previous infection is cleared at time *t*_clr_, immunity wanes at rate *w*.
(iii) In the additional risk model, the risk of an initial infection is determined as in the memoryless model (Equation 1), but the risk of subsequent infection is allowed to be higher. The force of infection thus includes a nonspecific additive risk factor, which is distinct from the effects of behavioral and demographic risk factors (Equation 4, Figure 2):

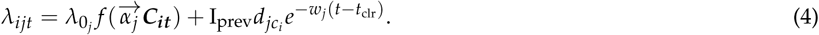

**Figure 2:**
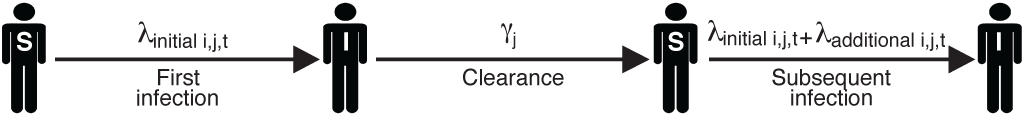
Schematic showing the dynamics for one individual host *i* and one HPV type *j*. The labels "S" and "I" indicate the susceptible and infected states, respectively. Before the initial infection, individual *i*’s per-capita, per-unit-time infection risk with type *j*, *λ*_*ijt*_, is determined by the type-specific baseline force of infection and the effect of host-specific risk factors, as in Equation 1. The duration of each infection is drawn from a Gamma distribution with mean 1/*γ*_*j*_. Previous infection modulates the risk of a subsequent infection as in Equation 4.

Here the additional risk parameter 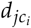 depends on the infecting type *j*, as well as the sexual subclass *c*_*i*_ of host *i*. The variable I_prev_ indicates whether there was a previous infection with type *j*.

We also included an observation model that determines the probability of the observed data using the known sensitivity and specificity of the HPV genotyping test. We fitted models to the data using a likelihood-based nonlinear fitting routine [49, 48]. To choose the model that best explains the data, we used the corrected Akaike Information Criterion (AICc), which balances the better fit of more complex models against the parsimony of simpler models, while including a correction for finite sample size [47]. The best model has the smallest AICc value.

We first tested whether infection with a particular HPV type reduces the risk of subsequent infection with the same type, allowing for variation in the degree and duration of protective homologous immunity (Equation 3). The homologous immunity model performed consistently worse than the memoryless model (Table 1), such that the additional parameters for homologous immunity increased model complexity without improving the fit, resulting in higher AICc scores (Table 1). There is therefore no evidence for homologous immunity against any of the six examined HPV types, suggesting that intra-type competition is weak to nonexistent.

**Table 1:** Comparison of Candidate Models

| Type | Model ($n$ parameters) | Log likelihood | SE | $\Delta\text{AICc}$ |
| --- | --- | --- | --- | --- |
| HPV62 | Memoryless (13) | -2026.1 | 1.4 | 389.6 |
|  | Homologous immunity (15) | -2027.4 | 1.5 | 396.3 |
|  | Additional risk (17) | -1827.2 | 0.4 | 0 |
| HPV16 | Memoryless (13) | -1663.2 | 1.1 | 117.9 |
|  | Homologous immunity (15) | -1663.7 | 0.7 | 123.1 |
|  | Additional risk (17) | -1600.1 | 0.1 | 0 |
| HPV89 | Memoryless (13) | -1902.3 | 1.3 | 137.4 |
|  | Homologous immunity (15) | -1902.2 | 1.5 | 141.2 |
|  | Additional risk (17) | -1829.5 | 0.1 | 0 |
| HPV51 | Memoryless (13) | -1670.8 | 0.6 | 63.9 |
|  | Homologous immunity (15) | -1670.4 | 1.7 | 67.3 |
|  | Additional risk (17) | -1634.7 | 0.7 | 0 |
| HPV84 | Memoryless (13) | -1979.6 | 1.7 | 270.3 |
|  | Homologous immunity (15) | -1980.5 | 1.1 | 276.3 |
|  | Additional risk (17) | -1840.3 | 0.4 | 0 |
| HPV6 | Memoryless (13) | -1461.0 | 1.0 | 121.7 |
|  | Homologous immunity (15) | -1461.5 | 1.3 | 126.8 |
|  | Additional risk (17) | -1396.1 | 0.6 | 0 |

Because separate types are less closely related, competition between types should be weaker still. Previous work has nevertheless speculated that there may be cross-immunity between HPV types. In particular, virus-like particles of HPV16 have been shown to induce a low level of neutralizing antibodies against HPV31, and clinical trial data suggest that current vaccines, which immunize against HPV16, provide partial protection against HPV31 [21, 55]. We therefore tested for competition between HPV16 and HPV31 by fitting a model in which previous infection with HPV31 affects the risk of infection with HPV16 (Supporting Information 2.2). Our estimate of the relative risk of HPV16 infection given previous HPV31 infection is centered around 1 (1.3, 95% C.I.: [0.5 , 2.0]), suggesting that the two types do not compete. This lack of an effect may be partly due to the low prevalence of HPV31 (Figure 1) and correspondingly low statistical power, but the lack of even a trend toward competition nevertheless suggests that there is no interaction. Accordingly, neither intra-type nor inter-type competition has strong effects on HPV dynamics.

### Past infection with HPV strongly increases the risk of future infection with the same type

The additional risk model fits the data vastly better than the homologous immunity and memoryless models (Table 1) for all 6 HPV types. The additional risk could reflect repeated exposure to the same HPV type by sexual activity with infected partners. To control for this possibility, we fitted the magnitude of the additional risk *d* separately for three different sexual subclasses: individuals reporting no recent sexual activity (celibate individuals), individuals reporting one recent sexual partner, and individuals reporting multiple recent sexual partners. In this analysis, we included only people who remained in the same sexual subclass for at least three consecutive years. For all six HPV types, previous infection significantly raises the risk of subsequent homologous infection irrespective of sexual subclass (Figure 3A).

**Figure 3:**
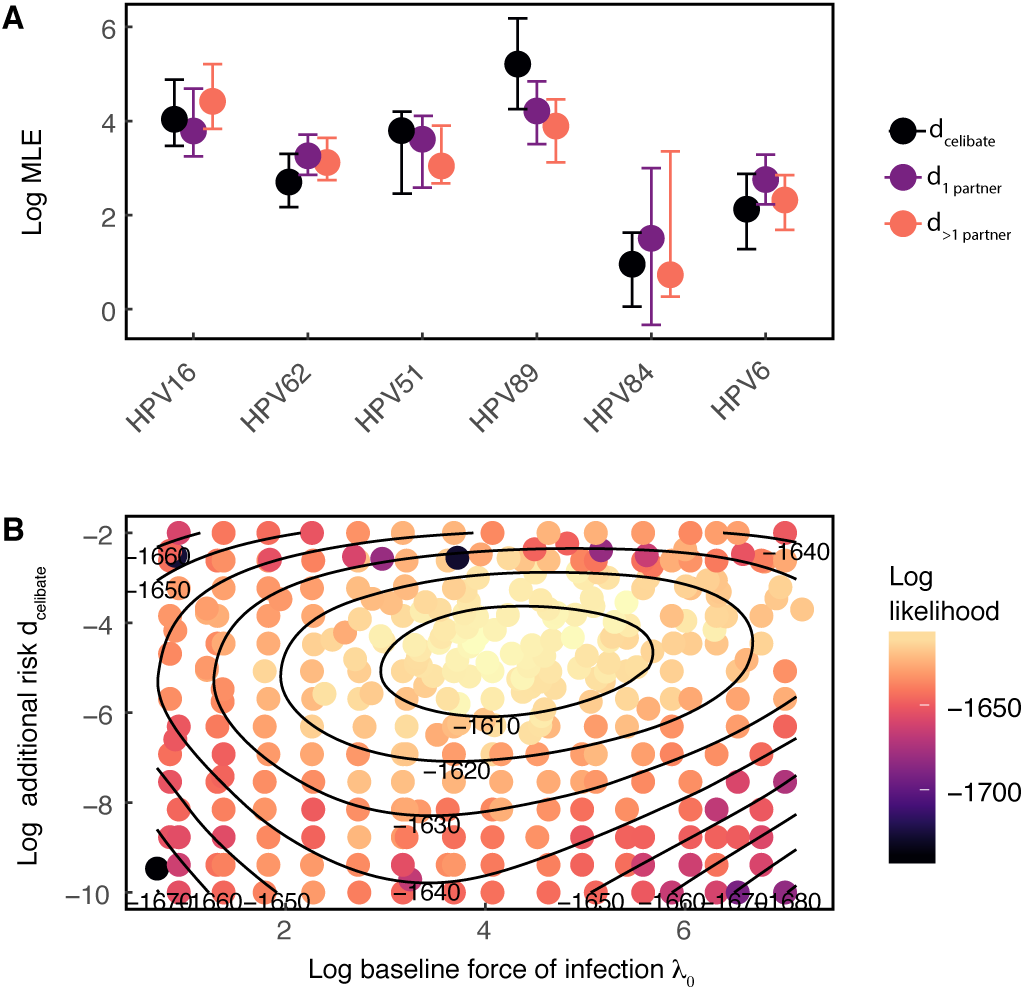
Inference of the additional risk *d* by sexual subclass. **A** Log estimates of *d* by sexual subclass for each HPV type. Points and error bars give the maximum likelihood estimate and 95% confidence interval, respectively. **B** Bivariate likelihood profile for the additional risk *d*_celibate_ and the baseline infection risk *λ*_0_ (shown on a log scale) Results shown for HPV16.

The high additional risk experienced by celibate individuals (*d*_celibate_) in particular strongly suggests that serial infections are driven by factors other than sexual transmission. Crucially, our estimate of the additional risk is uncorrelated with our estimate of baseline infection risk, *λ*_0*j*_ (Figure 3B), showing that the high estimated additional risk does not reflect statistical non-identifiability between additional risk and baseline risk. Estimates of the additional risk between different sexual subclasses have the same magnitude (Figure 3A), demonstrating that the additional risk varies only slightly across sexual subclasses. To quantify the impact of previous infection, we calculated the effect of previous infection on the total risk of a subsequent homologous infection at *t* = 0, 1, and 3 years after clearing the previous infection. For at least several years after the initial infection is cleared, the additional risk due to previous infection accounts for more than 90% of the force of infection (Figure S2). Moreover, immediately following a previous infection, the one-year probability of re-infection with HPV16 (Equation 5) is on average 20.4-fold higher than the probability of infection in a previously uninfected individual. The average increase is 19.1 - 20.5-fold among HPV types. At *t* = 3 years after clearing the previous infection, the one-year probability of reinfection with HPV16 remains 13.5-fold higher (7.4 - 20.5-fold among types).

The finding that initial infection strongly increases the risk of subsequent infection independently of sexual subclass is consistent with two major biological explanations: repeat infections, presumably due to auto-inoculation between anatomical sites, or episodic reactivation of latent infection. Because our models take into account 11 different risk factors besides previous infection, it is unlikely that the increased risk instead represents confounding by unmeasured host risk factors. Completely ruling out such unmeasured covariates is ultimately impossible without performing controlled experiments. As a preliminary approach to test for confounding, we repeated our estimation of the additional risk parameter *d* using a model that includes additional measured covariates for a total of 17 host-specific risk factors. In this more complex model, the additional risk *d* still accounts for more than 90% of the force of infection following clearance of the initial infection, and the model with more covariates did not provide a better fit to the data than the best-fit model with 11 covariates (Supporting Information 2.3). This result suggests that confounding by sexual risk factors is unlikely, while emphasizing the inability of traditional risk factors to explain the vast majority of HPV infections.

### Modest differences in host-specific risk factors suggest ecological differences between HPV types and highlight high-risk subpopulations

Although the additional risk conferred by past infection is substantial, a model without host-specific risk factors fits the data far worse than the full additional-risk model (Equation 4) for all types (Supporting Information 2.1). Moreover, the effects of these host-specific risk factors vary among HPV types. To understand the effects of this variation, we inserted our estimates of the baseline force of infection and the covariate effects for each HPV type into the best-fit model to calculate the distribution of infection risk in individuals who have never been infected. The expected time to infection (1/*λ*_*ijt*_), a measure of infection risk, is generally low but varies by orders of magnitude among individuals in the naive population (Figure S1), suggesting that the risk of initial infection with any type is concentrated among a few high-risk individuals.

Furthermore, the effects of each covariate show interesting similarities and differences between types (Figure 4). Increases in the number of recent female partners strongly increases risk for all HPV types, emphasizing the importance of heterosexual transmission. The addition of a single female sexual partner raises instantaneous infection risk by 80% to 120% among types compared to the risk in individuals with no recent female partners. The number of male partners, however, has divergent effects. The addition of a single male sexual partner *reduces* instantaneous risk with HPV16 by 50%, but *increases* the risk of infection with types HPV84 and HPV89 by 20% and 70%, respectively. Most other covariates were significant for only some of the types, indicating subtle differences between types. Correlations in risk factors between types were not associated with oncogenicity, such that HPV16 and HPV51, the two oncogenic types, were not more closely associated with one another based on their covariate profiles than they were to other types (Figure 4). Phylogenetic classification similarly did not reflect meaningful groupings of HPV types by their covariate profiles: HPV89, HPV62, and HPV84, all from the *α*-3 species, were not more closely associated to one another than to other types (Figure 4).

**Figure 4:**
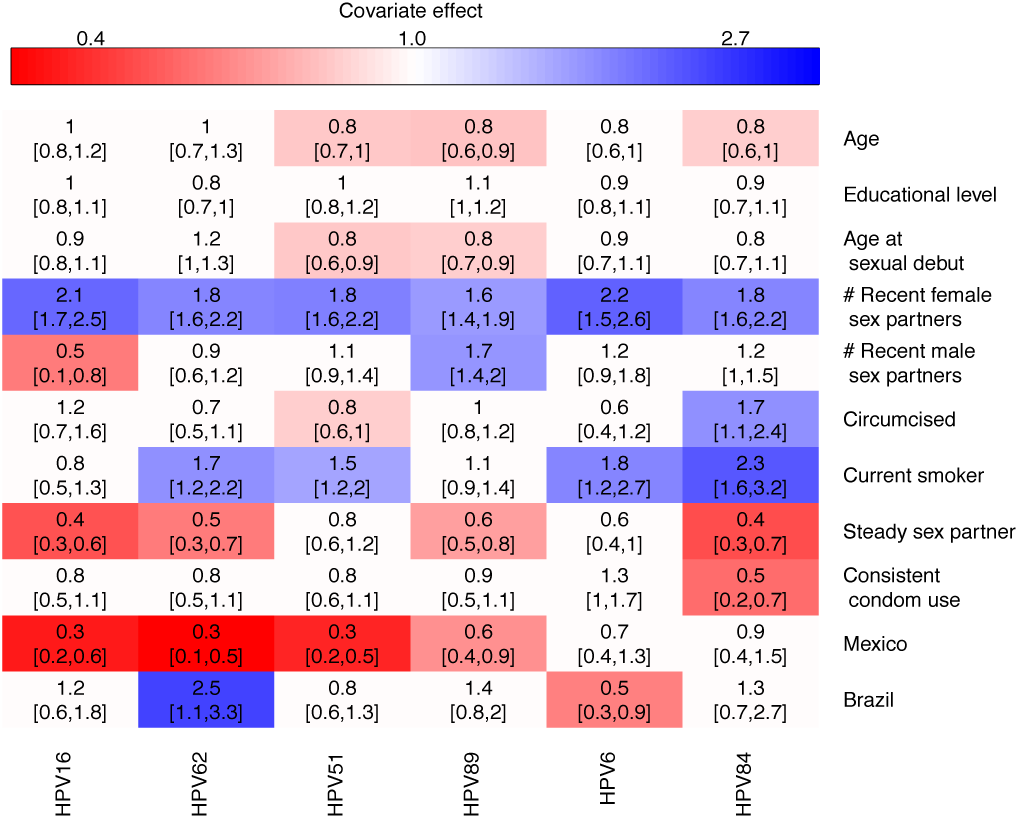
Inferred effect of host-specific covariates on the force of infection for each HPV type. Values provide the maximum likelihood parameter estimate and the 95% confidence interval. Cells colored in red denote statistically significant negative effects, and cells colored in blue denote statistically significant positive effects. HPV types are arranged by complete-linkage hierarchal clustering [63, 27].

Our estimates of the effects of type-specific risk factors are generally consistent with previous studies, but there are some exceptions. First, a previous analysis of the HIM data showed no difference in HPV16 incidence between men who have sex with men and men who have sex with women [85], but our results instead show that risk of HPV16 infection declines with an increase in the number of male sex partners, suggesting that HPV16 infection depends largely on heterosexual transmission. Meanwhile, our results show that the risk of infection with HPV51, HPV89, HPV6, and HPV84 increases with the number of male or female partners, consistent with previous work, although the effect was statistically significant only for HPV89. Finally, the previous analysis concluded that circumcision increases the risk of HPV51 [1], but our results suggest that circumcision instead reduces the risk of HPV51.

Several features of our modeling approach, however, may provide greater statistical power than previous analyses. First, our models account for variation in the risk per unit time, and they describe the dynamics in a way that is unconstrained and unbiased by the frequency of observations. Second, we infer the contribution of host-specific risk factors to the infection risk separately from the dynamics of previous infection. We therefore believe that our results are robust and that differences with respect to previous analyses arise from differences in the model structure.

## Discussion

Understanding the dynamics of HPV transmission is crucial to explain the coexistence of over 200 low-prevalence HPV types and to accurately predict the impact of multivalent HPV vaccines. We find little evidence of competition within or between HPV types, and instead we find high rates of reinfection or persistence by the same type. Measured epidemiological risk factors do explain at least some of the risk of initial infection, but they explain less than 10% of the risk after a man has recovered from his first infection. Previous infection with a particular type in contrast increases the risk of infection with the same type by 7-to 21-fold even three years after clearance. The inferred lack of both homologous and heterologous immunity implies that HPV type diversity cannot be explained by negative frequency-dependent selection, which promotes the coexistence of other immunologically distinct pathogen strains. The high prevalence of HPV instead results from continuous or repeated infection with multiple, apparently independent virus types. These types differ slightly in their risk factors for transmission to males, suggesting that ecological niche partitioning could play a modest role in further promoting coexistence.

Our results add to growing evidence that HPV infection, as opposed to vaccination, does not confer protective homologous immunity in men. Although we allowed the strength and duration of immunity to vary following initial infection, we could not detect any protection against reinfection with the same type. Because sterilizing immunity leads to lower overall infection rates, previous models that assume that infection results in protective immunity in men likely underestimated vaccine effectiveness [105, 24, 80, 7, 116, 11, 4, 26, 30, 61, 117, 5]. The well known effectiveness of the HPV vaccine may therefore be even higher than previously believed.

Our conclusion that there is no homologous immunity in men supports the hypothesis that genital infection differs between men and women. Although the durations and distributions of types in genital HPV infection are similar in men and women [44, 37], the prevalence of genital HPV is higher in men [44]. Acquired immunity has been proposed to explain declining cervical HPV prevalence by age in women in some countries [10, 102], whereas HPV prevalence does not change with age in men [34, 44]. One model showed that acquired immunity is necessary to explain age-specific patterns of prevalence in women [76], in contrast to our results in men. Furthermore, the seroprevalence of some types is higher in women than in men from the same source population [73, 17]. However, homologous protection in women is still likely to be weak [120, 6], supporting the idea that limited competition contributes to high HPV prevalence.

The high risk of recurring infection is consistent with either auto-inoculation across anatomic sites or with episodic reactivation of latent infection. The high risk of reinfection in celibate individuals makes confounding due to infected but unreported partners unlikely (Figure 3). The importance of auto-inoculation is supported by studies showing type-concordant HPV infection across anatomic sites. First, HPV DNA has been detected on the fingers of patients with genital warts [106], suggesting that HPV is transmitted between the hands and the genitals. Second, an analysis of a subcohort of the HIM study demonstrated a 1.5 to 15-fold increase in the risk of anal HPV infection after genital infection [85], and a similar analysis of sexually active women in Hawaii found a 20-fold increase in the risk of recurrent anal HPV infection given infection at the cervix [38]. Moreover, 63% of the cases of secondary anal HPV in the Hawaiian study occurred without a self-reported history of anal sex. Significantly, the magnitude of the effect in each of these studies is consistent with our estimate that previous infection leads to a 20-fold increase in the risk of subsequent infection.

Apparent reinfection could instead be due to the reactivation of an infection from a reservoir of latent virus. Whether HPV persists in a latent viral state remains unknown [100, 41], but an animal model of cottontail oral papillomavirus provides evidence of a latent viral reservoir in the basal layer of the mucosal epithelium [72]. HIV-positive women who are sexually inactive have a higher risk of recurrent HPV infection compared to HIV-negative controls [113], and among HIV-positive females, higher CD4+ T cell counts are negatively correlated with recurrent infection risk [110]. The latter two studies in particular have been interpreted as evidence of reactivation of latent infection [39], but either effect may instead result from increases in susceptibility to auto-infection rather than to diminished suppression of latent infection. Furthermore, the studies in HIV-positive individuals suggest that reactivation may require suppression of cellular immunity, and may therefore be rare in healthy individuals.

Vaccine efficacy trials furthermore suggest that reinfection is more common than reactivation. Women with previous HPV-related disease who received the quadrivalent vaccine were protected against HPV-related lesions [51], suggesting that reinfection is more common than reactivation. Similarly, one study showed 100% efficacy of the HPV vaccine against HPV-related disease in individuals with serological evidence of past HPV 6, 11, 16, or 18 infection [87]. Because HPV vaccines prevent infection by inducing antibodies that block viral entry at the epithelial basement membrane, such antibodies would likely not prevent reactivation of latent virus. If indeed vaccines do not affect the disease course of reactivated infections, then these vaccine efficacy trials suggest that the vaccine diminishes the incidence of HPV lesions by blocking true reinfection.

In short, although our estimate of the high additional risk from previous infection allows for the possibility that subsequent infections are due to the combined effects of reinfection and reactivation, other evidence suggests that autoinfection is more likely. This is important for vaccination policies, which often target people who are young enough that they are not yet sexually active. Vaccinating young people clearly reduces infection rates [74, 75], but if vaccination also reduces the risk of autoinfection, then vaccinating previously infected individuals may also strongly reduce HPV prevalence.

The type-specific effects of demographic and behavioral risk factors that we inferred suggest that modest differences exist in the host subpopulations supporting each HPV type. These differences highlight functional ecological distinctions between types that may further contribute to type coexistence For all types, the major determinant of infection risk is the number of recent female sex partners, suggesting a central role for heterosexual transmission. Additionally, current smoking increases risk in most types, consistent with other studies [33, 35, 44, 91]. Smoking can suppress mucosal and cellular immunity [62], but its effects may be confounded with other potentially unobserved, high-risk behaviors. Although shared risk factors account for most of the initial infection risk for each type, there are important distinctions too. For instance, the effect of circumcision on the risk of HPV infection is type-specific. Several randomized, controlled trials of male circumcision have demonstrated that circumcision protects against any HPV infection [3, 42], whereas observational studies have instead documented either no effect or increased risk [44, 68]. These conflicting results may have arisen from confounding by differences between types. Foreskin consists of unkeratinized mucosal epithelium, whereas the exposed tissue in circumcised men is made up of stratified squamous epithelium [42]. Our finding that circumcision increases the risk of infection with some HPV types (e.g. HPV84) and decreases the risk with others (e.g. HPV51) could reflect the adaptation of HPV types to different epithelia in the male genital tract.

Our modeling approach has several limitations, notably that we cannot distinguish between reactivation and auto-infection, and that we cannot rule out the possibility that unmeasured covariates affected our results. In addition, however, our model of homologous immunity assumes that protection arises immediately after infection, but protection may be lagged [80, 101]. The poor fit of the model that includes protective immunity nevertheless suggests that any such effects are quite weak, but allowing for more complex forms of homologous immunity is an important next step. Likewise, we assume that our estimate of the baseline infection risk for any type *j*, *λ*_0*j*_, captures the baseline risk of infection in naive hosts, but we have no information on HPV exposure or changes in risk factors that occurred before the study, nor do we have information on genetic differences in host susceptibility. Finally, the data track HPV infection in the genitals, ignoring other sites.

These caveats notwithstanding, the strong effect of initial infection on subsequent infection risk that we inferred is important for the design of epidemiological studies and models to inform public health policy. The effects of recurrent infection on subsequent risk suggest that vaccination may alter overall infection rates in ways that are not predicted from standard models of HPV dynamics [89, 24, 96]. The HPV vaccine therefore has the potential to lower HPV prevalence far more than previously expected.

## Materials and Methods

### Data

The HPV in Men (HIM) study is a multinational cohort study of HPV infection unvaccinated men aged 18-70 years. The HIM study provides the infection status with 37 types of HPV, confirmed by Linear Array PCR test [40], as well as host demographic and behavioral information. The study enrolled 4,123 participants between 2005 and 2009 and tracked men longitudinally over five years. Men were recruited from three cities: Tampa, Florida, USA; Cuernavaca, Mexico; and Sao Paulo, Brazil. Detailed study methods are described elsewhere [34, 35]. We excluded individuals that failed to meet the full eligibility criteria described by the HIM study [34], which included no prior diagnosis of genital cancer, warts, HIV, or other STIs. We classified men at each visit by sexual subclass according to their recent number of sexual partners, defined as the total number of reported sexual partners within the last six months. We defined three sexual subclasses: individuals that reported no sexual activity in the last six months (celibate individuals), individuals that reported one recent sex partner, and individuals that reported multiple (two or more) recent sex partners. The data were limited by individuals who switched between sexual subclasses. We included only the *n* =1,099 individuals that remained in their subclass for at least 3 years.

The data for each individual consist of binary time series describing infection status with each HPV type over a maximum of 10 clinic visits. The median number of visits among included participants was 10 visits. Data about host-specific covariates were drawn from a risk factor questionnaire administered at every visit. Additional characteristics of the data and a description of the covariate variables are provided in the Supporting Information (section 1).

### Model of HPV Dynamics

Our models are two-state discrete time partially observed Markov processes (POMPs), in which an individual is either infected (1) or uninfected (0) at any time with an HPV type. The 0 *→* 1 and 1 *→* 0 transitions are unobserved. The probability of infection (a 0 *→* 1 transition) is determined by the force of infection, *λ*. Infection of individual *i* with HPV type *j* occurs at rate *λ*_*ijt*_, and the time to infection is thus exponentially distributed. Thus, the probability of infection of individual *i* with type *j* (a 0*→*1 transition) over time interval Δ*t* is given by

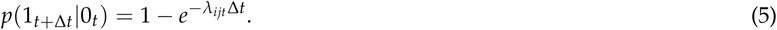

The measurement model is a Bernoulli distribution that relates each observation *Y*_*ijt*_ in an individual’s binary time series to the latent state *X*_*ijt*_ simulated under the model,

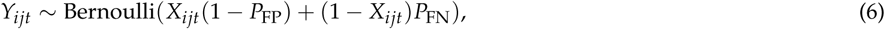

where *Y*_*ijt*_ is the observation of individual *i* with type *j* at time *t*, *X*_*ijt*_ is the corresponding latent state, *P*_FP_ is the rate of false positives, and *P*_FN_ is the rate of false negatives.

Table S3 describes the parameters for each candidate model. Each model was fit separately for each HPV type. Time-varying covariates (age, recent numbers of male and female sex partners, the presence of a steady sex partner, smoking status, educational level, and condom use) were updated at each visit. We parsimoniously used a log-linear model to model the covariate effects (Equation 2), such that each covariate effect *α* is directly interpretable as a multiplicative effect on the instantaneous risk of infection *λ*_*ijt*_. To identify the force of infection separately from the infection duration, the duration of infection with each HPV type *j* was drawn from a gamma distribution, where the shape (*k*_*j*_) and scale (*θ*_*j*_) were fixed to match the empirical distribution of infection durations in the dataset for type *j* (Table S4, Figure S7). The rate of false positives and false negatives in the measurement model were fixed based on the sensitivity and specificity of the genotyping test.

### Likelihood-based inference

Inference by maximum likelihood was carried out using multiple iterated filtering (MIF) [48] implemented in the R package “panelPomp” (version 0.3.1), which extends methods of the “pomp” package to longitudinal data [59, 58]. Briefly, iterated filtering is an algorithm that uses sequential Monte Carlo (SMC) to approximate maximum likelihood estimates of parameters from POMP models. SMC uses a population of particles drawn from the parameters of a given model to generate Monte Carlo samples of the latent dynamic variables and evaluate the likelihood of observed time series [2, 20]. Iterated filtering successively filters the particle population, perturbing the parameters between iterations. The perturbations decrease in amplitude over time, allowing convergence at the maximum likelihood estimate.

The extension of SMC and iterated filtering to longitudinal panel data has been previously described [97], and we extended longitudinal POMP methods to binary data. The data for each HPV type is a set of binary time series, or panel units, describing the observed infection trajectory for each individual. The panel POMP contains a POMP model for each individual, and individuals share parameters. To evaluate the likelihood of a shared parameter set, SMC is carried out over the time series for each individual to generate a unit log-likelihood. The log likelihood of the panel POMP object is the sum of the individuals’ log likelihoods. All optimization routines were carried out using 20,000 particles to overcome high Monte Carlo error (Supporting Information 3).

For each model, we initialized the iterated filtering with 100 random parameter combinations. Optimization involved series of successive MIF searches, with the output of each search serving as the initial conditions for the subsequent search. The likelihood of the output for each search was calculated by averaging the likelihood from ten passes through the particle filter using 50,000 particles. The optimization was repeated until additional operations did not arrive at a higher maximum likelihood (Supporting Information 3).

For model selection, we used the corrected Akaike Information Criterion (AICc) [47]. We obtained parameter estimates (maximum likelihood estimates and 95% confidence intervals) by constructing likelihood profiles. We used Monte Carlo Adjusted Profile methods [49] to obtain a smoothed estimate of the profile that accounts for the increased Monte Carlo error associated with longitudinal data. The lower and upper limits of the 95% confidence interval were the points that lay 1.92 log-likelihood units below the maximum likelihood estimate on the smoothed profile curve (the points corresponding to one-half the 95% critical value for a *χ*^2^ distribution with one degree of freedom).

## Data accessibility

The data and code for this analysis are available at https://github.com/cobeylab/HPV-model.

## Author Contributions

S.R., S.C., and G.D. designed the research; S.R. performed research and analysis; E.B. provided computational expertise; V.D. provided statistical expertise; A.G., L.V., and E.L. provided data; A.G. provided clinical expertise; S.R., S.C., and G.D. wrote the manuscript.

## Acknowledgements

Ken Alexander was instrumental in the initiation of this project. We thank Daniel Zinder for helpful discussions about the model construction. This work was completed with resources provided by the University of Chicago Research Computing Center and with funding provided by the National Institute of Allergies and Infectious Diseases (NIH F30AI124636 to S.R.) and by the University of Chicago Medical Scientist Training Program (NIH T32GM007281).

## Supporting Information

**Figure S1:**
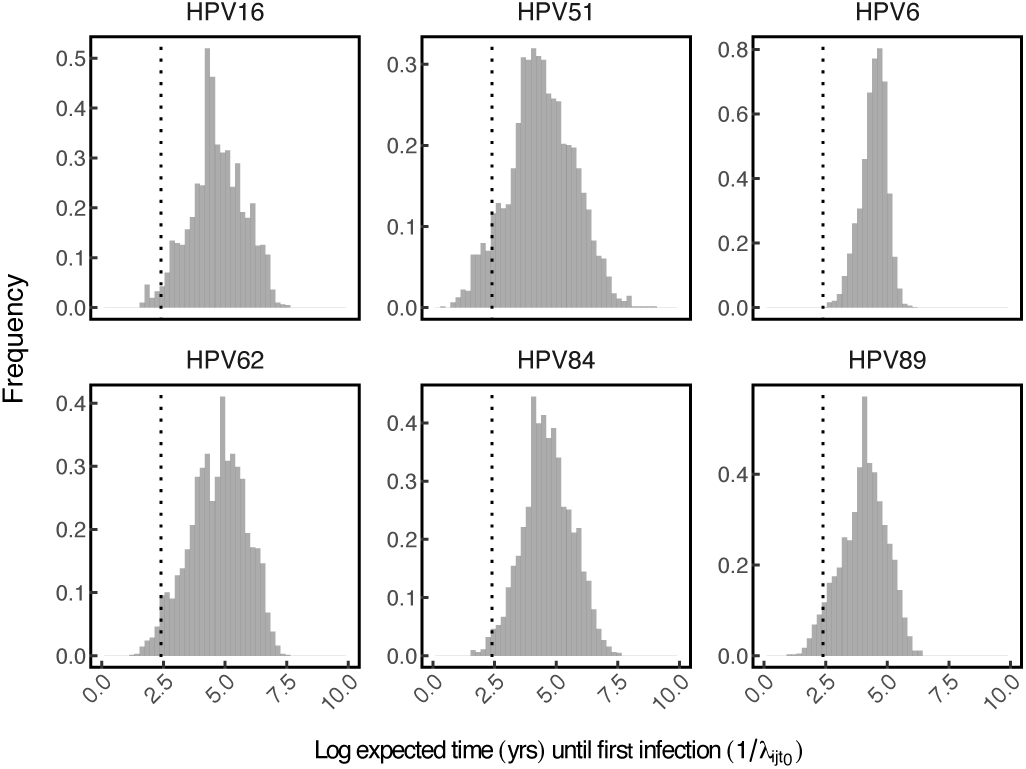
Distribution of the (log) baseline expected time to infection (*t*_expected_ = 1/*λ*_*ij*_) in uninfected individuals assuming no prior infection, such that 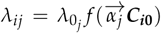. For each individual, *λ*_*ij*_ was calculated using the maximum likelihood estimate for each element in 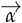 and the individual-specific covariates ***C*_*i*_0**, which were reported at the baseline visit (*t* = 0). The y axis reports frequency, while the vertical dashed line in each panel marks an expected time to infection of 10 years. Thus, only the portion of the distribution to the left of the dashed line in each panel represents individuals for whom the the expected infection time falls within the next ten years.

**Figure S2:**
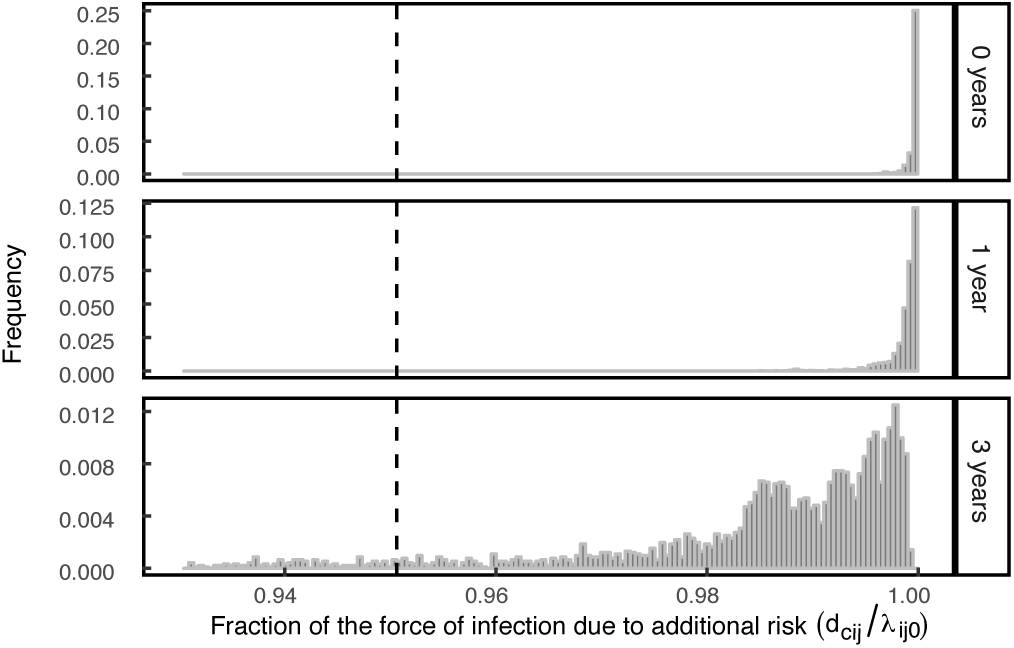
Contribution of the effect of previous infection to overall infection risk in individuals with respect to HPV16. Histograms show the distribution of the fraction of the overall force of infection *λ*_*ijt*_ made up by the additional risk *d* in the population at various times post-clearance of the precedent infection. The mean value for the population-level distribution at each time point is given by the vertical dotted line.

**Figure S3:**
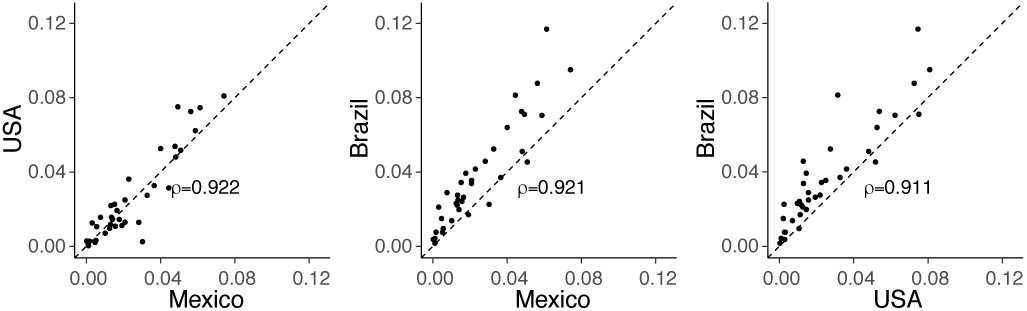
Comparison of the time-averaged prevalence of HPV types in the HIM Dataset. The black dotted lines indicates 1:1 correlation, and *ρ* denotes Pearson’s correlation coefficient (p *<* 10*-*15 with 35 degrees of freedom for each comparison).

**Figure S4:**
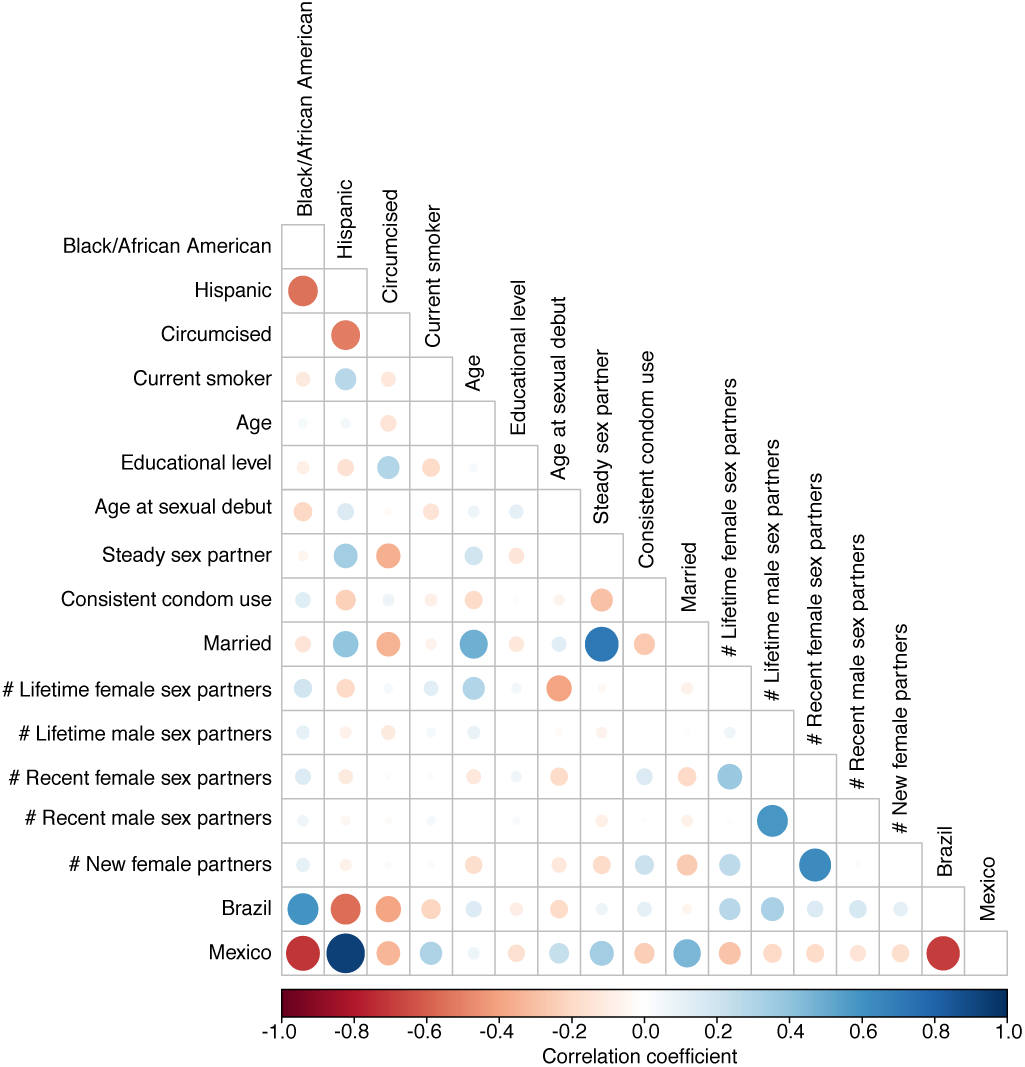
Correlations between the full candidate set of self-reported covariates at the baseline visit. Pearson product-moment correlations were calculated between continuous variables, polyserial correlations (inferred latent correlations between continuous and categorical variables) were calculated between numeric and binary variables, and polychoric correlations (inferred latent correlations between categorical variables) were calculated between binary variables [22, 88]. All correlations shown were significant at the *α* = .05 significance level based on tests for bivariate normality.

**Figure S5:**
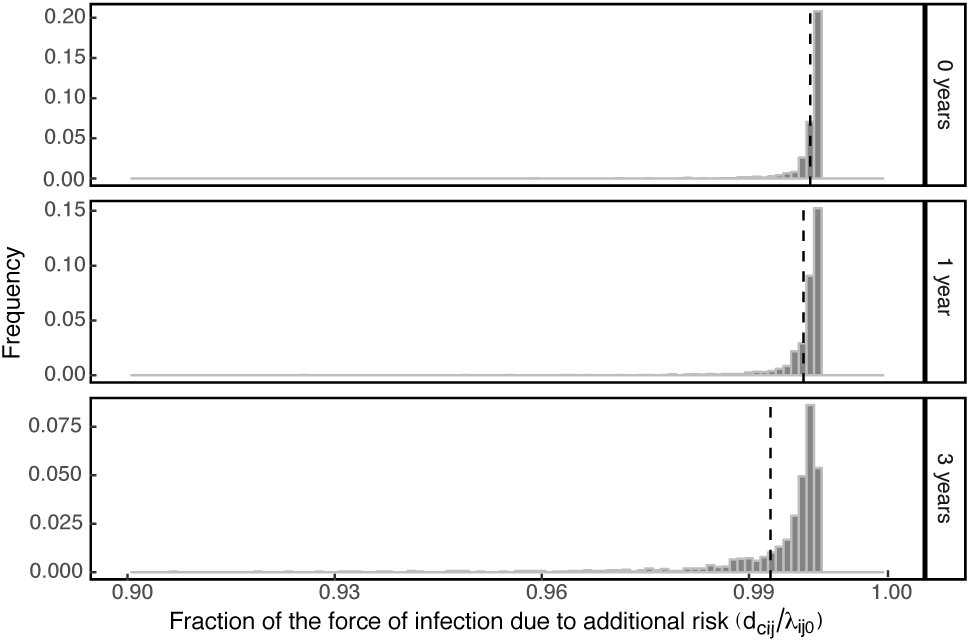
Contribution of the effect of previous infection to the overall infection risk among individuals for HPV16, using a model with 17 covariates. Histograms show the distribution of the fraction of the overall force of infection *λ*_*ijt*_ made up by the additional risk d in the population at various times post-clearance of the precedent infection. The mean value for each distribution is given by the vertical dotted line.

**Figure S6:**
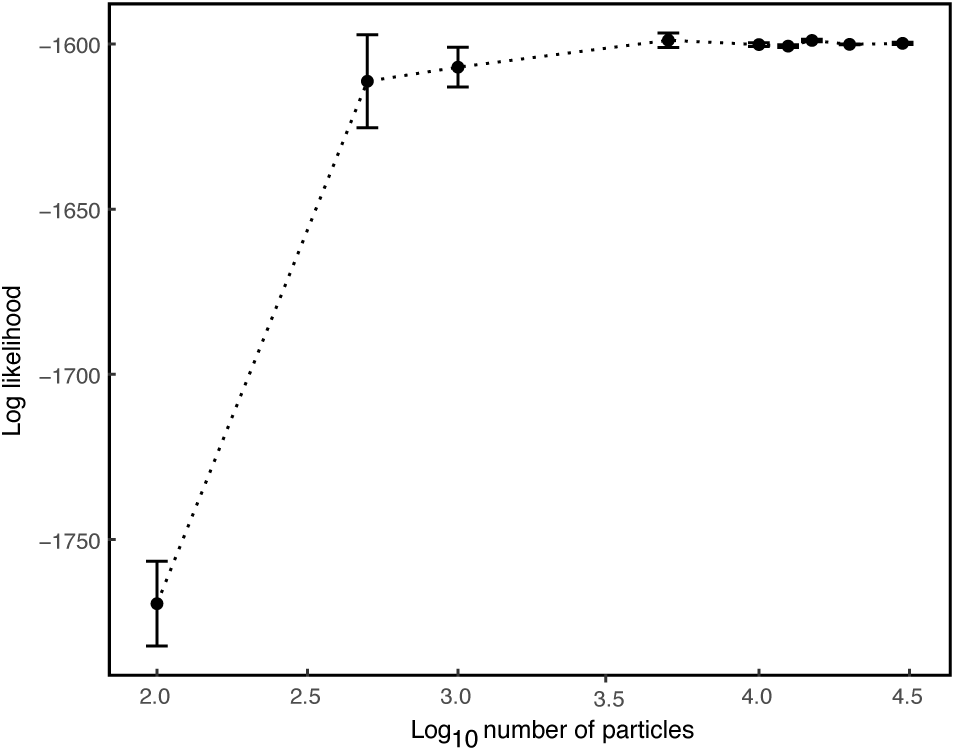
Likelihood of the maximum likelihood parameter set for HPV16 calculated at increasing particle sizes. Point estimates and error bars represent the mean and standard error, respectively, of 10 particle filter replicates.

**Figure S7:**
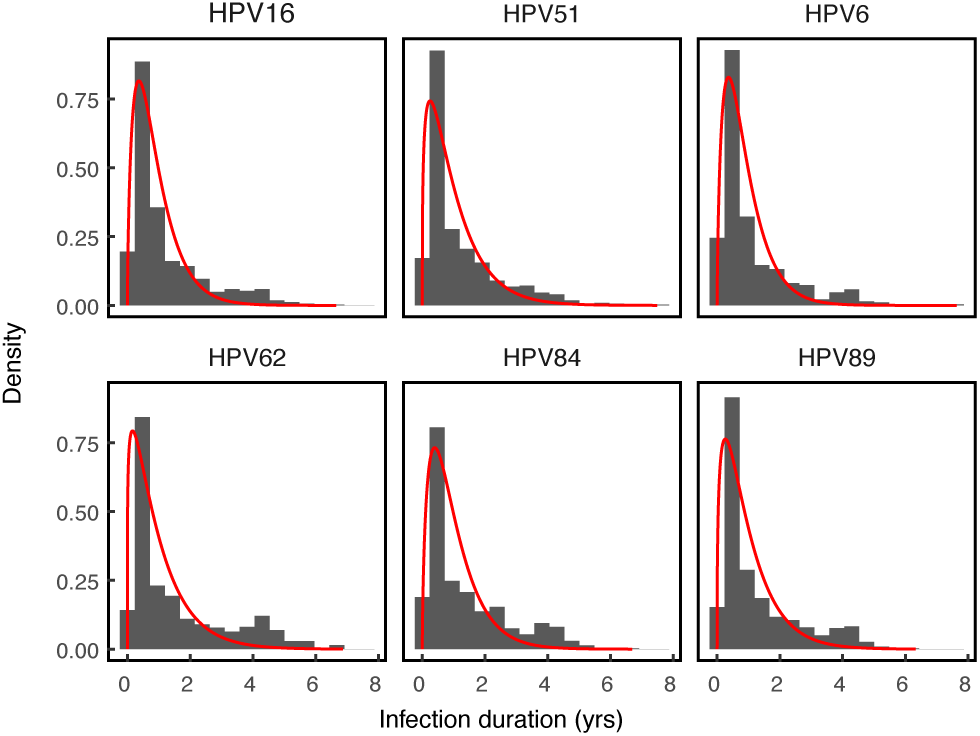
Empirical distribution of infection durations for each HPV type (grey histogram) with an overlying gamma distribution having the same mean and coefficient of variation as in the data (red).

## 1 Details of the HPV in Men data and selection of covariate variables

We identified 3,656 eligible participants from the 4,123 men enrolled in the HIM study as of October 2014. The eligibility criteria, taken from Giuliano et al. [34], included no prior diagnosis of genital cancer, warts, HIV, or other STIs. Of the eligible participants, we excluded *n* = 575 participants that had missing data at the time of enrollment for the 11 covariates that we included in our three candidate models. We then divided the remaining 3,081 participants into three sexual subclasses based on the number of recent sex partners. For all covariates, "recent" activity indicates activity in the past six months, as reported at each clinic visit. The three sexual subclasses were individuals reporting no recent sexual activity and no recent sex partners (celibate individuals), individuals with one recent sex partner, and individuals with two or more recent sex partners. To account for the effects of sexual subclass on the infection risk, we restricted our analysis to include only the *n* = 1, 099 individuals who remained in the same subclass for at least three years over the course of their participation in the study.

Covariates were derived from a risk factor questionnaire that was administered at each visit, which was described and validated previously [34, 86, 69, 91, 23, 1]. Questions covered sociodemographic characteristics, sexual behaviors, sex partnerships, and condom use. Participants were asked to recall their recent behavior, such as recent number of male or female sexual partners, where "recent" referred to behavior over the past six months. Participants had the option of refusing to answer any question, and refusals were treated as missing values as in Giuliano et al. [34], such that a missing covariate was assigned its value at the closest visit.

Covariates were selected for inclusion in the model based on known risk factors identified in the literature for HPV in men [34, 33, 35, 37, 83, 84, 85, 94, 44, 91, 23, 69, 1]. We also included country of residence as a covariate. The full set of candidate covariates (Figure S4) included race (black/African American or other), ethnicity (Hispanic or other), age, age at sexual debut, lifetime numbers of male and female sexual partners, numbers of recent male and female sexual partners, numbers of new male and female sexual partners, presence of a steady sexual partner, marital status, level of education, circumcision status (confirmed by a clinician at the baseline visit), whether or not the participant was a current smoker, whether or not the participant used condoms for the majority of recent sexual encounters, Brazilian nationality, and Mexican nationality, with US nationality used as the baseline. To decrease statistical non-identifiability and to increase computational feasibility, we reduced the full candidate set to a subset of covariates based on observed correlations (Figure S4). Among highly correlated pairs of similar covariates, we heuristically selected one representative. For example, we chose to include recent numbers of sexual partners instead of lifetime numbers of sexual partners and the presence of a steady sexual partner instead of marital status. The variables describing race and ethnicity were strongly correlated with country of origin, so we excluded them. Table S1 gives the final subset of covariates included in the model.

**Table S1:** Covariates included in the analyses.

|  | Name | Structure | Notes |
| --- | --- | --- | --- |
| 1 | Age | Continuous |  |
| 2 | Educational Level | Continuous | Levels: <12 yrs (0), 12 yrs (1), 13-15 yrs (2), 16 yrs (3), >= 17 yrs (4) |
| 3 | Age at sexual debut | Continuous |  |
| 4 | Recent female sexual partners | Continuous | 0,1,2,>=3 different partners within the last six months |
| 5 | Recent male sexual partners | Continuous | 0,1,2,>=3 different partners within the last six months |
| 6 | Circumcision status | Binary |  |
| 7 | Smoking status | Binary | Current smoker |
| 8 | Steady sex partner | Binary | Presence of a steady sexual partner |
| 9 | Consistent condom use | Binary | Use of a condom more than 50% of the time during intercourse |
| 10 | Brazil | Binary | Country of residence |
| 11 | Mexico | Binary | Country of residence |

## 2 Alternative and additional models

### 2.1 Additional risk only model

In this model, the force of infection *λ*_*ijt*_ for individual *i* with type *j* at time *t* was determined only by the baseline infection risk for type *j* and by the effect of previous infection, so that the behavioral and demographic risk factors had no effect. The force of infection is then:

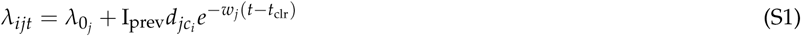

For each HPV type, this model out-performed the memoryless model but it performed much worse than the model that also took into account the covariates (Table S2).

**Table S2:** Performance of the additional risk only model for each HPV type relative to the full additional risk model.

| Type | Log likelihood | SE | $\Delta\text{AICc}$ |
| --- | --- | --- | --- |
| HPV16 | -1,636.5 | 0.6 | 50.6 |
| HPV51 | -1,651.5 | 1.8 | 11.7 |
| HPV6 | -1,403.6 | 0.3 | 8.3 |
| HPV62 | -1,877.0 | 0.4 | 66.8 |
| HPV84 | -1,870.0 | 0.1 | 37.3 |
| HPV89 | -1,843.4 | 0.8 | 22.5 |

### 2.2 HPV16/HPV31 interaction model

To test whether infection with HPV31 affects the risk of infection with HPV16, we fitted a model in which the force of infection of HPV16 depends on whether an individual has ever been infected with HPV31. In this model, we included a covariate variable I_HPV31_, updated at each visit, that indicated whether individual *i* was currently or previously infected with HPV31. The force of infection *λ*_*ijt*_ was thus:

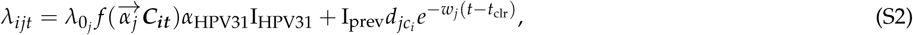

where *α*_HPV31_ gives the effect of previous or current infection with HPV31 on the risk of infection with HPV16. Our goal was to estimate the direction and strength of interaction between HPV16 and HPV31. Our estimate of the interaction parameter was centered around 1 (MLE 1.3, CI[0.5,2.0]), indicating no significant interaction.

### 2.3 Additional Covariates model

We added additional covariates to the best-fit model for HPV16, using the full set of 17 original candidate covariates (Figure S4) in the additional risk model. Our goal was to estimate the contribution of the additional risk to the force of infection when allowing for a large number of host risk factors. We used the parameters from the maximum likelihood estimate of the model to estimate the fraction of the force of infection made up by the additional risk *d* among individuals. This model involved inference for *n*=1019 individuals that had complete covariate information at baseline for the full set of covariates. To compare this model to the best-fit (additional risk) model, we fitted the additional risk model for the same *n*=1019 individuals and obtained the maximum likelihood. We find that the additional risk *d* still accounts on average for over 90% of the risk for several years after infection clearance (Figure S5). The maximum likelihood and AICc for the model with additional covariates were -1468.4 (SE .63) and 2983.9, respectively, whereas the maximum log likelihood and AICc for the model without additional covariates, fitted to the data from the same individuals, were -1472.8 (SE .86) and 2979.6, respectively. Thus, the ΔAICc for the additional covariates model is 4.3, showing that the improvement in the likelihood from the additional complexity of the additional covariates does not provide a better explanation of the data than the additional risk model.

## 3 Monte Carlo error in inference from binary panel data

Advances in simulation-based Monte Carlo methods have made it possible to fit complex models to large datasets. We took advantage of extensions of multiple iterated filtering (MIF) [48] to the case of panel data. Iterated filtering uses sequential Monte Carlo, also known as particle filtering, to estimate the likelihood of partially observed Markov process (POMP) models. Sequential Monte Carlo uses stochastic simulations of dynamical models to produce successive populations of weighted particles. Each particle represents a Monte Carlo sample from the probability density of the latent process, conditional on the parameters and the previous observations. As the particle population is propagated along the time series, the particles are weighted and resampled at each data point, and the likelihood of each observation is estimated as the weighted average of the particles.

Large data sets and complex models can result in non-negligible Monte Carlo error in estimated likelihoods. The structure of panel data, a collection of time series that are dynamically independent apart from shared model parameters, yields high Monte Carlo error that often makes it infeasible to calculate the likelihood with an error of less than one log likelihood unit [49]. This is important because a standard approach to calculating 95% confidence intervals relies on the observation that parameter values with log likelihood scores that are within 1.92 units of the maximum log likelihood fall within the 95% confidence interval [9]. Because high levels of Monte Carlo error can make it difficult to accurately estimate likelihoods, high Monte Carlo error rates also make it difficult to estimate 95% confidence intervals. Ionides et al. [49] show that one solution to this problem is to approximate the likelihood in the region of the maximum likelihood by fitting a quadratic to to likelihood scores from a large sample of parameter values, which can in turn be used to directly estimate the confidence bounds. Ionides et al. validated this approach with panel data [49], and here we apply it to the case of binary panel data.

To test our approach, before we carried out the model fitting, we quantified the effect of particle size on the likelihood for a given set of parameters. As is often the case in simulation-based approaches, the Monte Carlo error in our simulations is high enough that most particles have very low likelihoods. As a result, our likelihood estimates at first improve rapidly with increases in the number of particles (Figure S6). For particle numbers above about 5000, however, further increases in particle numbers have at most weak effects. We therefore used 20,000 particles for the iterated filtering and 50,000 particles for evaluations of the likelihood by particle filtering. We also accounted for Monte Carlo error in maximum likelihood estimation by initiating a large number of independent MIF searches (*n* = 100) at random parameter values for any given model. Each of the 100 searches began with 200 MIF iterations (.75 cooling factor) and was continued successively (100 iterations, .75 cooling factor) until the maximum likelihood for that particular search was stationary within one log likelihood unit. To identify the MLE for a given model, we required that three searches independently arrive within two log likelihood units of the maximum likelihood value.

## 4 Model parameters

**Table S3:** Description of model parameters

| Model | Parameter type | Parameter name | Fixed/estimated/nuisance |
| --- | --- | --- | --- |
| All Models | Baseline infection risk | $\lambda_{0j}$ | Estimated |
| All Models | Infection duration | Shape of gamma distribution ($k_j$) | Fixed |
| All Models | Infection duration | Scale of gamma distribution ($\theta_j$) | Fixed |
| All models | Covariate effects ($\alpha$) | Age | Estimated |
| All Models | Covariate effects ($\alpha$) | Educational status | Estimated |
| All Models | Covariate effects ($\alpha$) | Age at sexual debut | Estimated |
| All Models | Covariate effects ($\alpha$) | Circumcision status | Estimated |
| All Models | Covariate effects ($\alpha$) | Number of recent female sexual partners | Estimated |
| All Models | Covariate effects ($\alpha$) | Number of recent male sexual partners | Estimated |
| All Models | Covariate effects ($\alpha$) | Steady sexual partner | Estimated |
| All Models | Covariate effects ($\alpha$) | Consistent condom use | Estimated |
| All Models | Covariate effects ($\alpha$) | Current smoking status | Estimated |
| All Models | Covariate effects ($\alpha$) | Mexican nationality ($\alpha_{MX}$) | Estimated |
| All Models | Covariate effects ($\alpha$) | Brazilian nationality ($\alpha_{BZ}$) | Estimated |
| All Models | Initial conditions | Probability of initial infection ($P_{\text{initial},j}$) | Estimated |
| All Models | Initial conditions | Duration of remaining infection ($F_{\text{remaining}}$) | Nuisance |
| All Models | Initial conditions | Probability of previous infection ($p_{\text{past}}$) | Nuisance |
| All Models | Measurement model | False positive rate ($P_{FP}$) | Fixed |
| All Models | Measurement model | False negative rate ($P_{FN}$) | Fixed |
| Homologous immunity | Immunity dynamics | Magnitude of scaled risk ($d$) | Estimated |
| Homologous immunity | Immunity dynamics | Rate of waning of immunity ($w$) | Estimated |
| Additional risk model | Additional risk ($d$) | $d_{\text{celibate}}$ | Estimated |
| Additional risk model | Additional risk ($d$) | $d_{1\text{partner}}$ | Estimated |
| Additional risk model | Additional risk ($d$) | $d_{\text{multiple}}$ | Estimated |

### Fixed parameters

The false positive and false negative rates of HPV detection were fixed according to the high sensitivity (96%) and specificity (99%) of the Roche Linear Array HPV genotyping test reported by the manufacturer (Roche Diagnostics), which has been confirmed by other analyses [109, 40, 77]. The duration of each simulated infection with each HPV type *j* was drawn from a γ(*k*_*j*_, *θ*_*j*_) distribution, where *k*_*j*_ and *θ*_*j*_ were fixed according to the empirical distribution of infection durations in the data for type *j* (Figure S7). Following Giuliano et al. [34], we required two consecutive negative visits following a positive visit for any HPV type to constitute an observed clearance. Thus, we treated 1-0-1 transitions as false negatives, thereby adjusting data a priori based on the assumption that infection in such cases was actually continuous. Our justification was two-fold: first, this approach allowed us to account for false negatives in HPV sampling beyond the laboratory specifications of the HPV test that were included in the observation model. Second, the approach ensured that the high rates of reinfection estimated by the best-fit model were not the result of failing to account for false clearances, ensuring in turn that our estimates of reinfection rates would be conservative.

**Table S4:** Values of the parameters describing infection durations

| Type | $k_j$ | $\theta_j$ |
| --- | --- | --- |
| HPV62 | 1.20 | 0.81 |
| HPV84 | 1.67 | 0.59 |
| HPV89 | 1.37 | 0.70 |
| HPV16 | 1.71 | 0.52 |
| HPV51 | 1.33 | 0.75 |
| HPV6 | 1.71 | 0.51 |

### Nuisance parameters

Two nuisance parameters were used to generate the initial conditions for model simulations but were not estimated. For individuals that were initially infected during any simulation, the fraction of an infection duration that they had experienced prior to time *t* = 0 was given by the nuisance parameter *F*_remaining_, which was drawn from a uniform (0,1) distribution for each model realization. For individuals that were initially uninfected during any simulation, the probability that the individual had previously been infected at some point in time was given by *p*_past_, which was drawn from a uniform (0,1) distribution for each model realization.

### Estimated parameters for the best-fit model

**Table S5:**
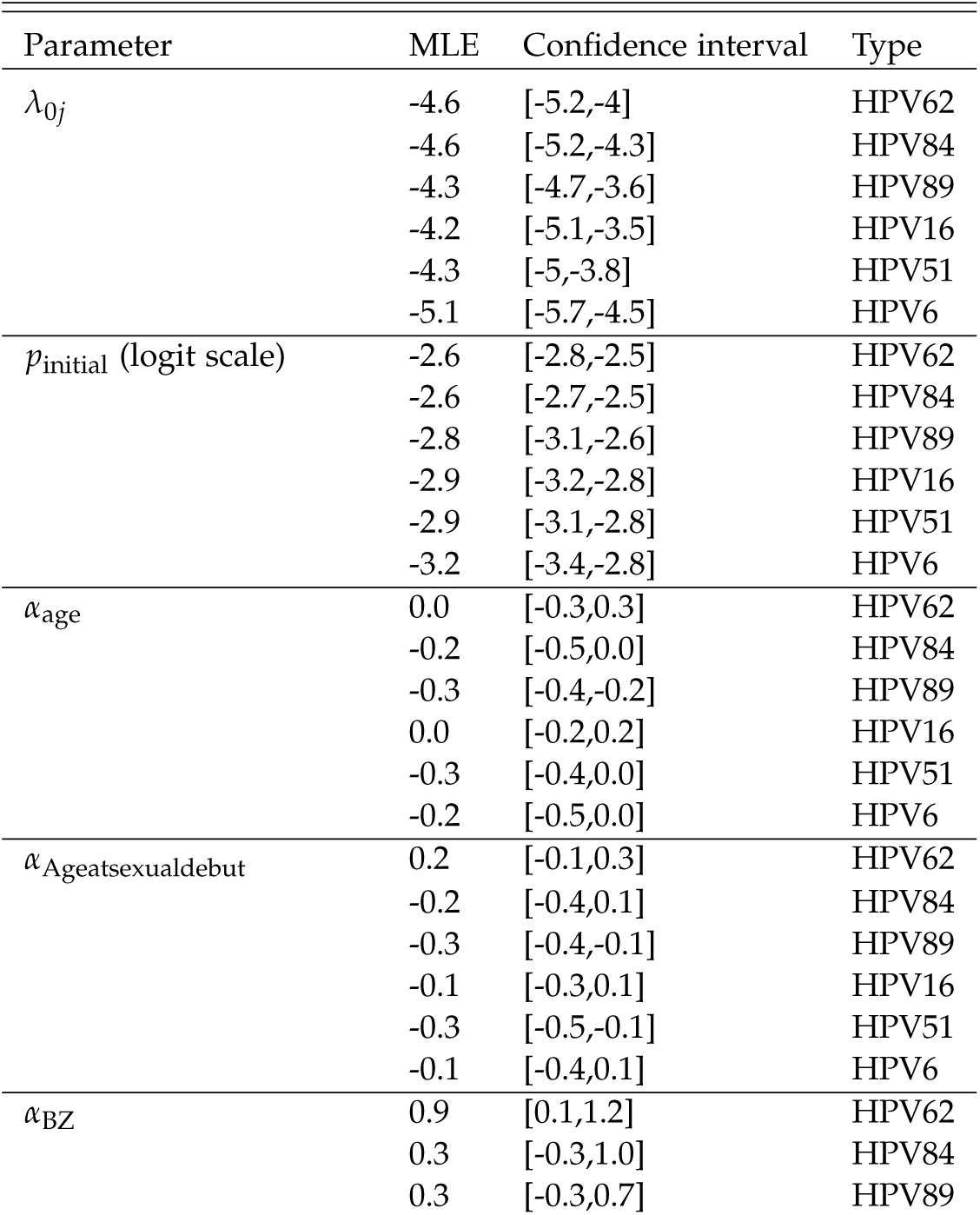
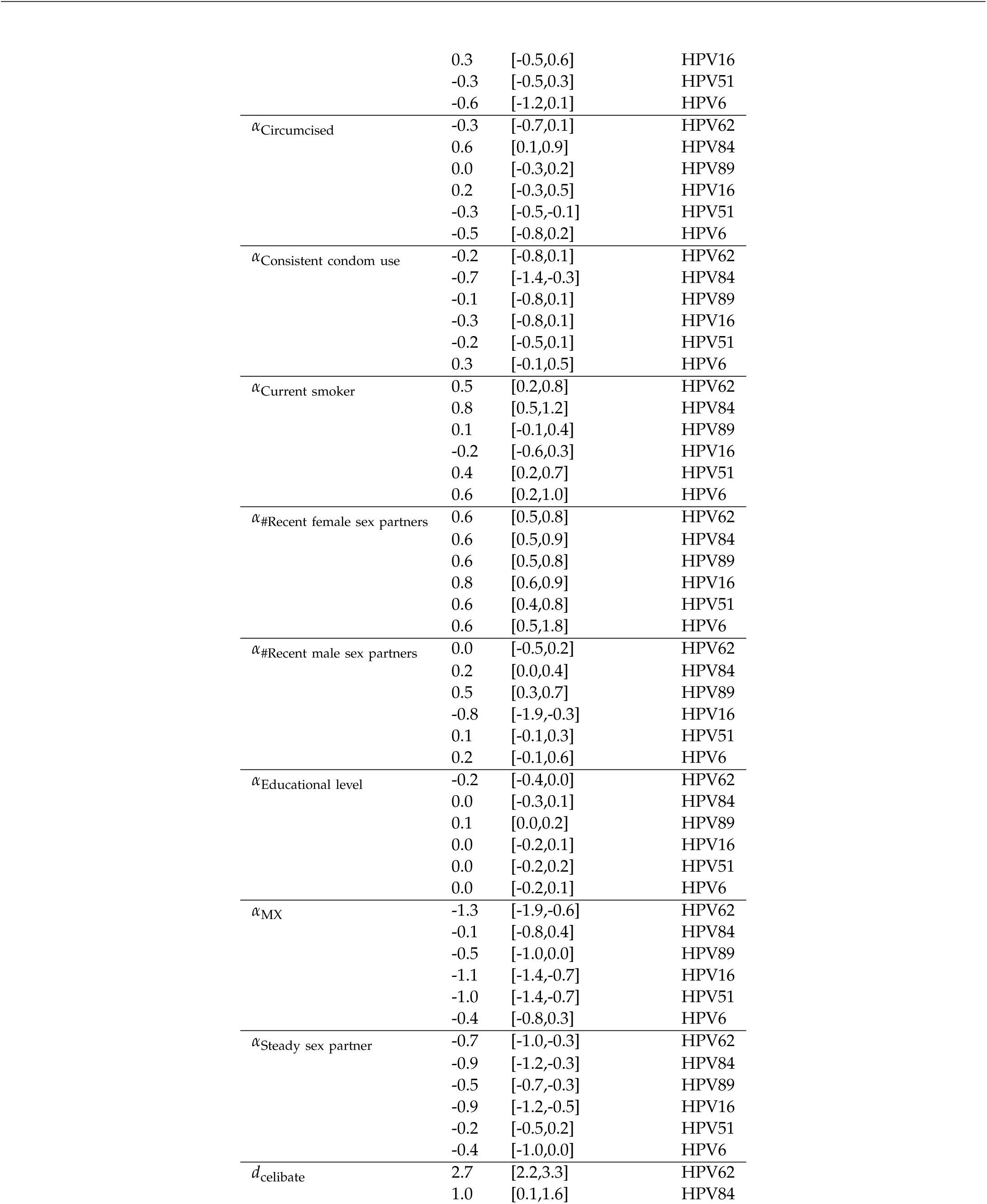
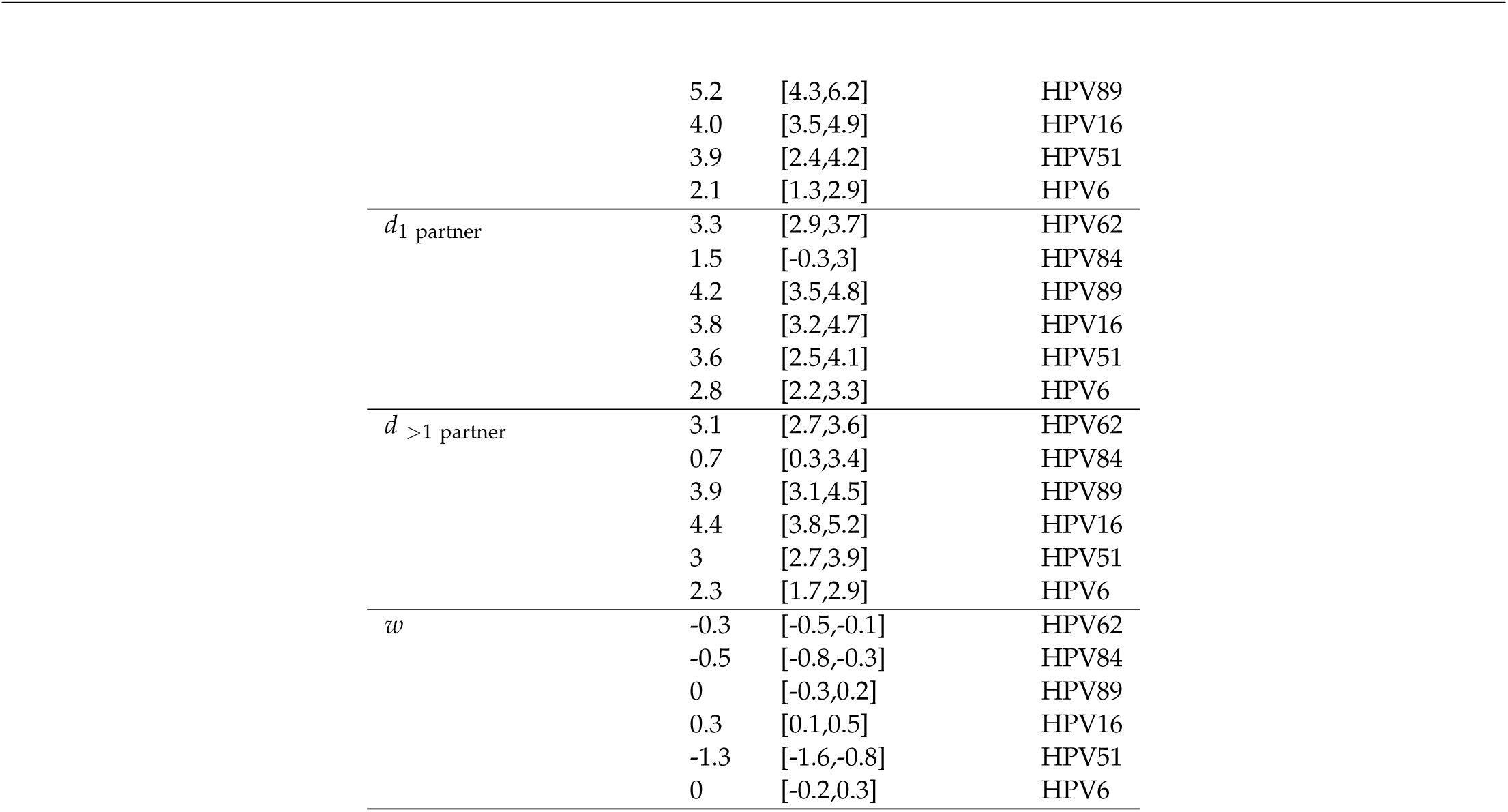
Values of the estimated parameters. Estimates are provided on a log scale unless otherwise indicated.

| Parameter | MLE | Confidence interval | Type |
| --- | --- | --- | --- |
| $\lambda_{0j}$ | -4.6 | [-5.2,-4] | HPV62 |
|  | -4.6 | [-5.2,-4.3] | HPV84 |
|  | -4.3 | [-4.7,-3.6] | HPV89 |
|  | -4.2 | [-5.1,-3.5] | HPV16 |
|  | -4.3 | [-5,-3.8] | HPV51 |
|  | -5.1 | [-5.7,-4.5] | HPV6 |
| $p_{\text{initial}}$ (logit scale) | -2.6 | [-2.8,-2.5] | HPV62 |
|  | -2.6 | [-2.7,-2.5] | HPV84 |
|  | -2.8 | [-3.1,-2.6] | HPV89 |
|  | -2.9 | [-3.2,-2.8] | HPV16 |
|  | -2.9 | [-3.1,-2.8] | HPV51 |
|  | -3.2 | [-3.4,-2.8] | HPV6 |
| $\alpha_{\text{age}}$ | 0.0 | [-0.3,0.3] | HPV62 |
|  | -0.2 | [-0.5,0.0] | HPV84 |
|  | -0.3 | [-0.4,-0.2] | HPV89 |
|  | 0.0 | [-0.2,0.2] | HPV16 |
|  | -0.3 | [-0.4,0.0] | HPV51 |
|  | -0.2 | [-0.5,0.0] | HPV6 |
| $\alpha_{\text{Ageatsexualdebut}}$ | 0.2 | [-0.1,0.3] | HPV62 |
|  | -0.2 | [-0.4,0.1] | HPV84 |
|  | -0.3 | [-0.4,-0.1] | HPV89 |
|  | -0.1 | [-0.3,0.1] | HPV16 |
|  | -0.3 | [-0.5,-0.1] | HPV51 |
|  | -0.1 | [-0.4,0.1] | HPV6 |
| $\alpha_{\text{BZ}}$ | 0.9 | [0.1,1.2] | HPV62 |
|  | 0.3 | [-0.3,1.0] | HPV84 |
|  | 0.3 | [-0.3,0.7] | HPV89 |
|  | 0.3 | [-0.5,0.6] | HPV16 |
|  | -0.3 | [-0.5,0.3] | HPV51 |
|  | -0.6 | [-1.2,0.1] | HPV6 |
| $\alpha_{\text{Circumcised}}$ | -0.3 | [-0.7,0.1] | HPV62 |
|  | 0.6 | [0.1,0.9] | HPV84 |
|  | 0.0 | [-0.3,0.2] | HPV89 |
|  | 0.2 | [-0.3,0.5] | HPV16 |
|  | -0.3 | [-0.5,-0.1] | HPV51 |
|  | -0.5 | [-0.8,0.2] | HPV6 |
| $\alpha_{\text{Consistent condom use}}$ | -0.2 | [-0.8,0.1] | HPV62 |
|  | -0.7 | [-1.4,-0.3] | HPV84 |
|  | -0.1 | [-0.8,0.1] | HPV89 |
|  | -0.3 | [-0.8,0.1] | HPV16 |
|  | -0.2 | [-0.5,0.1] | HPV51 |
|  | 0.3 | [-0.1,0.5] | HPV6 |
| $\alpha_{\text{Current smoker}}$ | 0.5 | [0.2,0.8] | HPV62 |
|  | 0.8 | [0.5,1.2] | HPV84 |
|  | 0.1 | [-0.1,0.4] | HPV89 |
|  | -0.2 | [-0.6,0.3] | HPV16 |
|  | 0.4 | [0.2,0.7] | HPV51 |
|  | 0.6 | [0.2,1.0] | HPV6 |
| $\alpha_{\text{\#Recent female sex partners}}$ | 0.6 | [0.5,0.8] | HPV62 |
|  | 0.6 | [0.5,0.9] | HPV84 |
|  | 0.6 | [0.5,0.8] | HPV89 |
|  | 0.8 | [0.6,0.9] | HPV16 |
|  | 0.6 | [0.4,0.8] | HPV51 |
|  | 0.6 | [0.5,1.8] | HPV6 |
| $\alpha_{\text{\#Recent male sex partners}}$ | 0.0 | [-0.5,0.2] | HPV62 |
|  | 0.2 | [0.0,0.4] | HPV84 |
|  | 0.5 | [0.3,0.7] | HPV89 |
|  | -0.8 | [-1.9,-0.3] | HPV16 |
|  | 0.1 | [-0.1,0.3] | HPV51 |
|  | 0.2 | [-0.1,0.6] | HPV6 |
| $\alpha_{\text{Educational level}}$ | -0.2 | [-0.4,0.0] | HPV62 |
|  | 0.0 | [-0.3,0.1] | HPV84 |
|  | 0.1 | [0.0,0.2] | HPV89 |
|  | 0.0 | [-0.2,0.1] | HPV16 |
|  | 0.0 | [-0.2,0.2] | HPV51 |
|  | 0.0 | [-0.2,0.1] | HPV6 |
| $\alpha_{\text{MX}}$ | -1.3 | [-1.9,-0.6] | HPV62 |
|  | -0.1 | [-0.8,0.4] | HPV84 |
|  | -0.5 | [-1.0,0.0] | HPV89 |
|  | -1.1 | [-1.4,-0.7] | HPV16 |
|  | -1.0 | [-1.4,-0.7] | HPV51 |
|  | -0.4 | [-0.8,0.3] | HPV6 |
| $\alpha_{\text{Steady sex partner}}$ | -0.7 | [-1.0,-0.3] | HPV62 |
|  | -0.9 | [-1.2,-0.3] | HPV84 |
|  | -0.5 | [-0.7,-0.3] | HPV89 |
|  | -0.9 | [-1.2,-0.5] | HPV16 |
|  | -0.2 | [-0.5,0.2] | HPV51 |
|  | -0.4 | [-1.0,0.0] | HPV6 |
| $d_{\text{celibate}}$ | 2.7 | [2.2,3.3] | HPV62 |
|  | 1.0 | [0.1,1.6] | HPV84 |
|  | 5.2 | [4.3,6.2] | HPV89 |
|  | 4.0 | [3.5,4.9] | HPV16 |
|  | 3.9 | [2.4,4.2] | HPV51 |
|  | 2.1 | [1.3,2.9] | HPV6 |
| $d_{1 \text{ partner}}$ | 3.3 | [2.9,3.7] | HPV62 |
|  | 1.5 | [-0.3,3] | HPV84 |
|  | 4.2 | [3.5,4.8] | HPV89 |
|  | 3.8 | [3.2,4.7] | HPV16 |
|  | 3.6 | [2.5,4.1] | HPV51 |
|  | 2.8 | [2.2,3.3] | HPV6 |
| $d_{>1 \text{ partner}}$ | 3.1 | [2.7,3.6] | HPV62 |
|  | 0.7 | [0.3,3.4] | HPV84 |
|  | 3.9 | [3.1,4.5] | HPV89 |
|  | 4.4 | [3.8,5.2] | HPV16 |
|  | 3 | [2.7,3.9] | HPV51 |
|  | 2.3 | [1.7,2.9] | HPV6 |
| $w$ | -0.3 | [-0.5,-0.1] | HPV62 |
|  | -0.5 | [-0.8,-0.3] | HPV84 |
|  | 0 | [-0.3,0.2] | HPV89 |
|  | 0.3 | [0.1,0.5] | HPV16 |
|  | -1.3 | [-1.6,-0.8] | HPV51 |
|  | 0 | [-0.2,0.3] | HPV6 |

**Figure S8:**
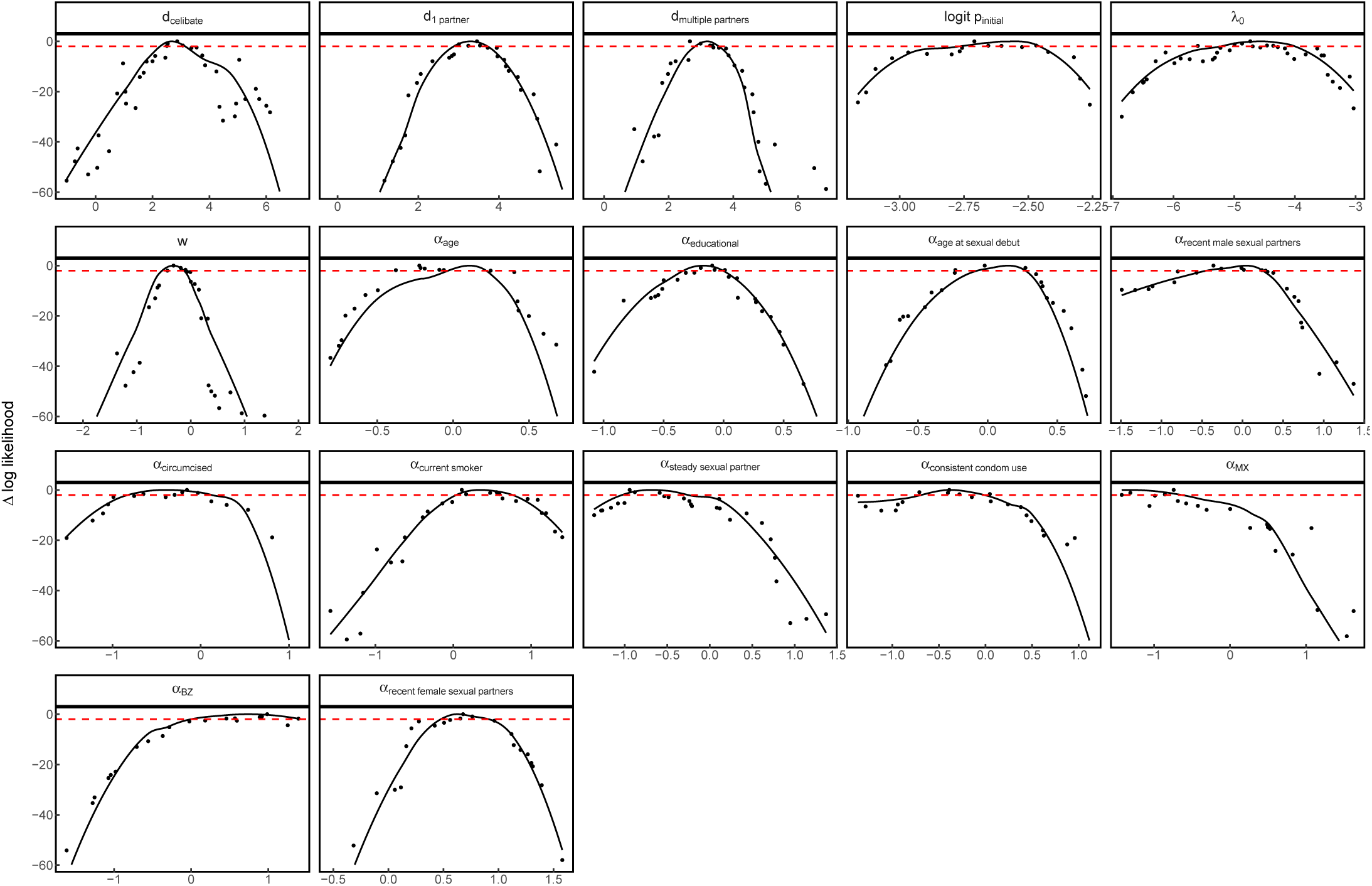
HPV62: Likelihood profiles for each estimated parameter. The x axis gives the log of the profile parameter value unless the logit distribution is specified. The y axis gives the likelihood relative to the maximum likelihood for the additional risk model for HPV62. The red dashed horizontal line indicates the cutoff of 1.92 log likelihood units used to determine the confidence interval.

**Figure S9:**
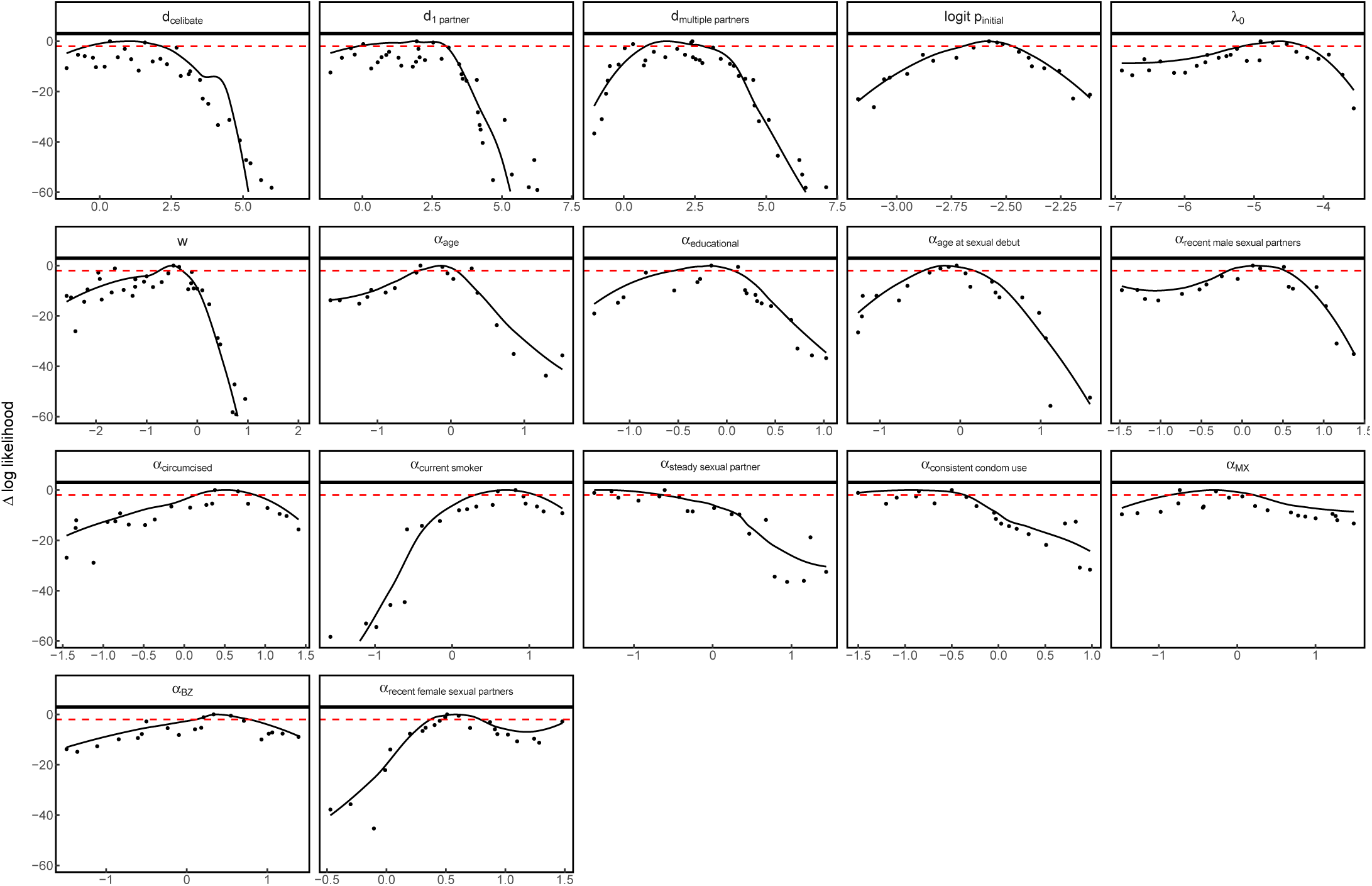
HPV84: Likelihood profiles for each estimated parameter, as in Figure S8.

**Figure S10:**
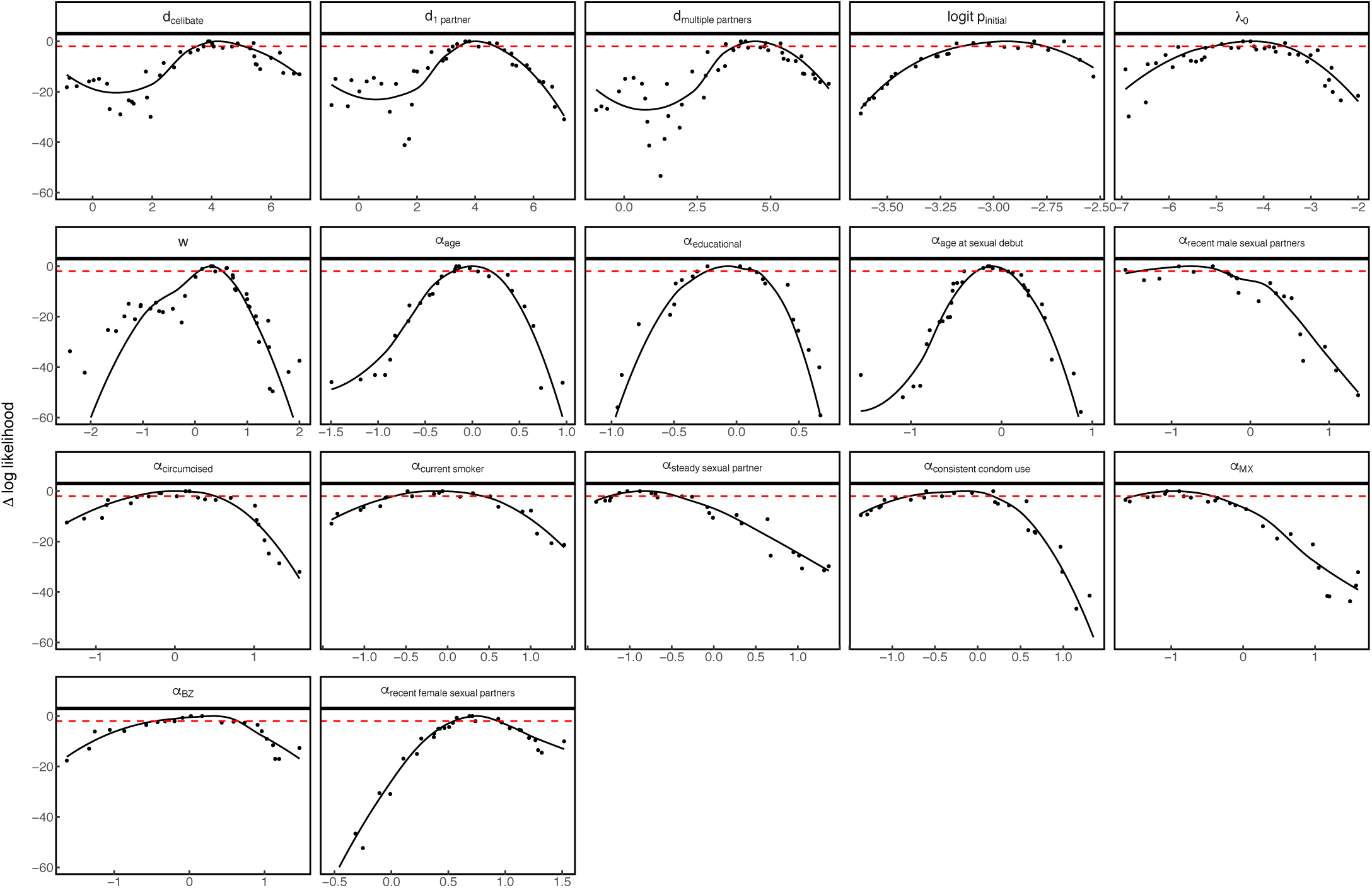
HPV16: Likelihood profiles for each estimated parameter, as in Figure S8.

**Figure S11:**
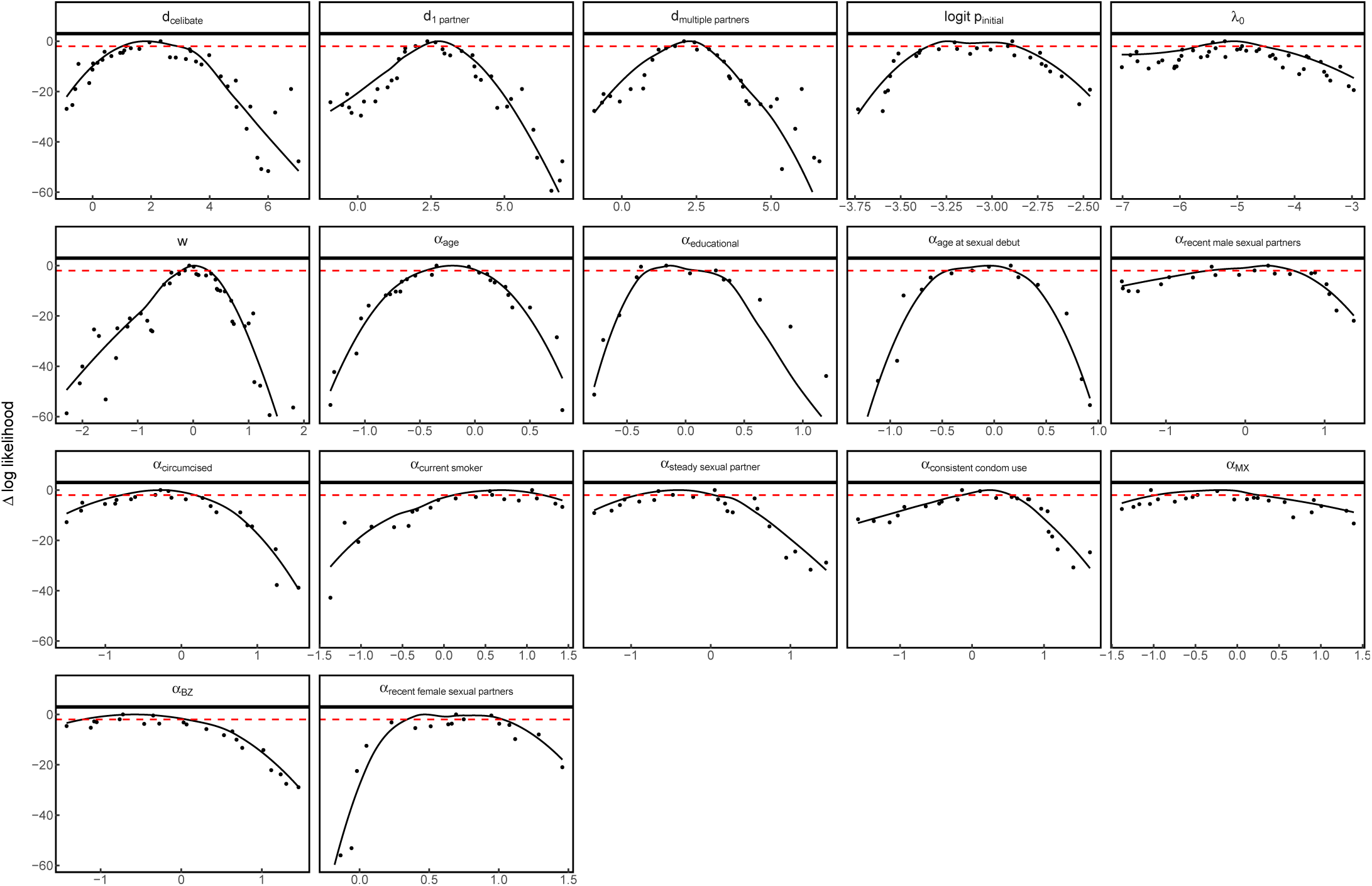
HPV6:Likelihood profiles for each estimated parameter, as in Figure S8.

**Figure S12:**
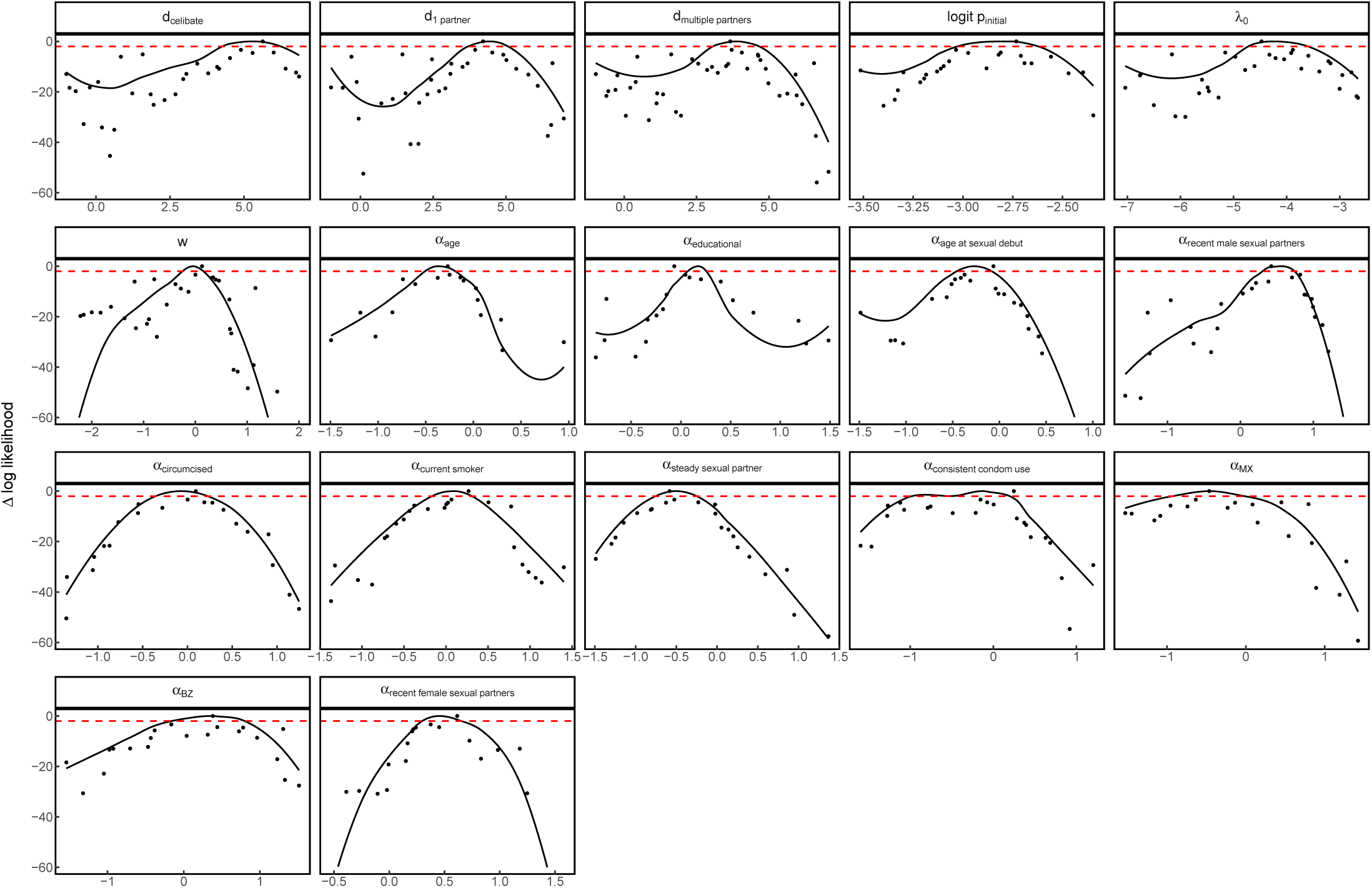
HPV89: Likelihood profiles for each estimated parameter, as in Figure S8.

**Figure S13:**
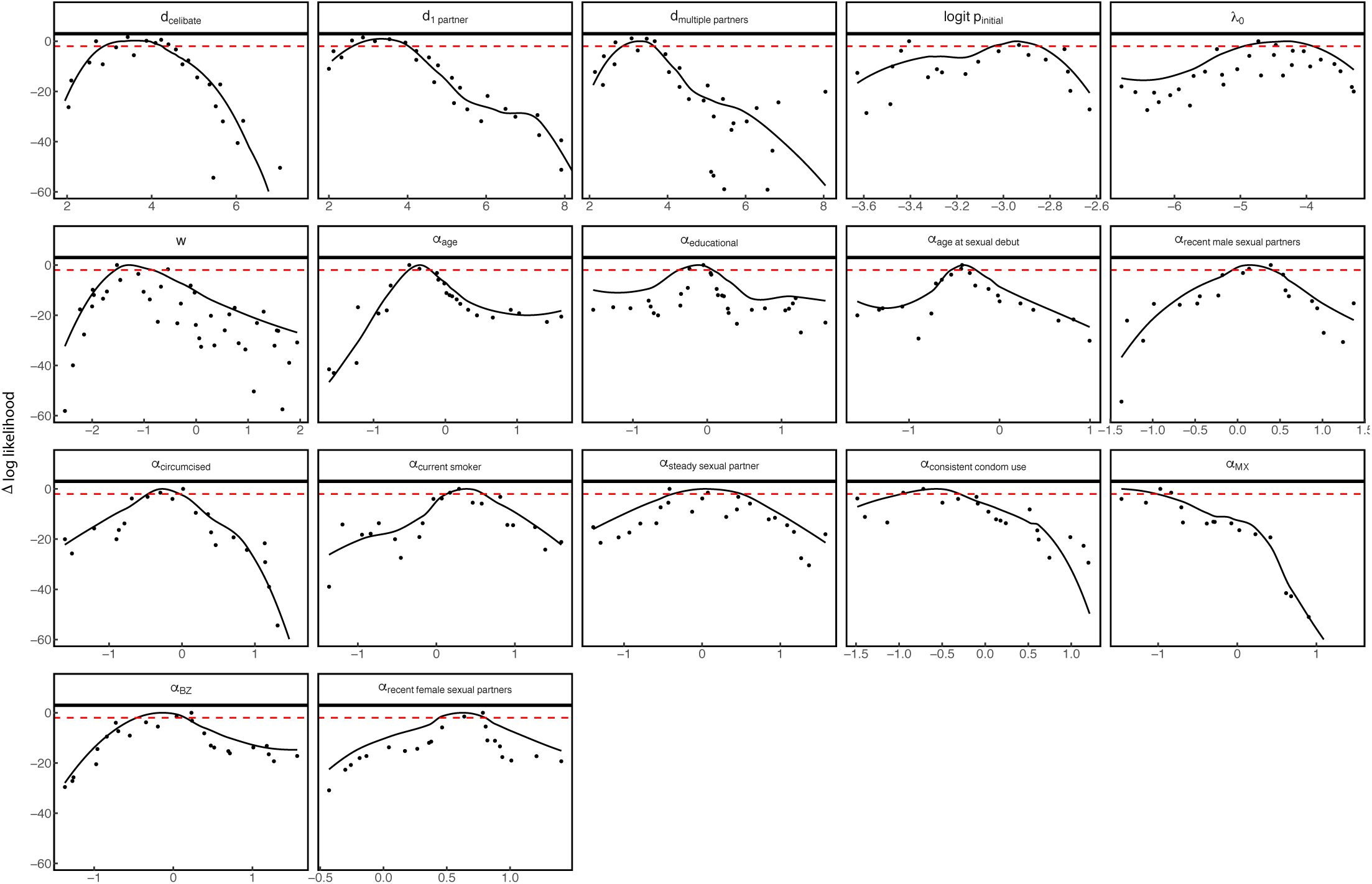
HPV51: Likelihood profiles for each estimated parameter, as in Figure S8.

